# Pupil self-regulation modulates markers of cortical excitability and cortical arousal

**DOI:** 10.1101/2024.09.04.611153

**Authors:** Marieke Lieve Weijs, Silvia Missura, Weronika Potok-Szybińska, Marc Bächinger, Bianca Badii, Manuel Carro Dominguez, Nicole Wenderoth, Sarah Nadine Meissner

**Author notes:** authors contributed equally (Marieke Lieve Weijs, Silvia Missura, Weronika Potok-Szybińska, Nicole Wenderoth, Sarah Nadine Meissner). Address for correspondence: Marieke Lieve Weijs or Nicole Wenderoth, Neural Control of Movement Laboratory, Department of Health Sciences and Technology, ETH Zurich, Gloriastrasse 37/39, 8092 Zurich, Switzerland.

## Abstract

The brain’s arousal state (i.e., central arousal) is regulated by multiple neuromodulatory nuclei in the brainstem and significantly influences high-level cognitive processes. By exploiting the mechanistic connection between the locus coeruleus (LC), a key regulator of central arousal, and pupil dynamics, we recently demonstrated that participants can gain volitional control over arousal-regulating centers including the LC using a pupil-based biofeedback approach. Here, we test whether pupil-based biofeedback modulates electrophysiological markers of cortical excitability, cortical arousal, and phasic LC activity. Combining pupil-based biofeedback with single-pulse TMS, EEG recordings, and an auditory oddball task revealed three main results: pupil self-regulation significantly modulates (i) cortical excitability, (ii) the EEG spectral slope, a marker of cortical arousal, and (iii) the P300 response to target tones, an event-related potential suggested to be tightly linked to phasic LC activity. Interestingly, pupil self-regulation strength was linearly linked to the modulation of the spectral slope, suggesting a common physiological mechanism. Here, we have shown that pupil-based biofeedback modulates fundamental aspects of brain function. Whether this method could further be used to modulate these aspects in case of disturbances associated with neurological and psychiatric disorders needs to be investigated in future studies.

## Introduction

The arousal state of the brain (i.e., central arousal) influences various ongoing high-level cognitive processes and is closely related to phenomena such as attention, anxiety or the sleep-wake cycle^1,2^. Central arousal is controlled by multiple neuromodulatory nuclei, including the noradrenergic (NA) locus coeruleus (LC), as well as dopaminergic and cholinergic regions^3–5^. Particularly, the LC has been shown to be involved in arousal regulation^3,6,7^ and to substantially modulate neural processing via diverse NA projections to cortical and subcortical brain regions^7,8^. Under constant lighting conditions, changes in the eye’s pupil size have been linked directly or indirectly to neuromodulatory systems involved in arousal regulation^9,10^ with the strongest evidence found for the LC-NA system^10–13^.

We have recently inverted this mechanism and demonstrated that participants can learn to volitionally control pupillary dynamics when trained with a pupil-based biofeedback approach (pupil-BF)^14^: By using arousing or relaxing mental strategies during real-time pupil-BF, participants were able to learn to volitionally increase (i.e., upregulate) and decrease (i.e., downregulate) their pupil size. Crucially, by combining pupil size up- and downregulation using pupil-BF and spatially rich fMRI measurements, our analyses showed that such self-regulation is related to systematic activity changes in brainstem regions involved in arousal and autonomic regulation with the most robust modulation observed in the LC. Additionally, pupil self-regulation affected dopaminergic and cholinergic brain regions^14^. Furthermore, pupil self-regulation modulated heart rate (HR) and heart rate variability (HRV)^14^, as predicted by existing LC projections to cardiovascular control centers^15^. However, it is currently unknown whether self-regulating pupil diameter as a proxy of LC-NA activity, modulates the arousal state of cortical areas.

Cortical arousal has been measured using different neurophysiological markers. First, numerous studies have provided causal evidence of noradrenergic contributions to changes in cortical excitability, as measured by transcranial magnetic stimulation (TMS) induced motor evoked potentials (MEPs). Specifically, administrating pharmacological NA application with TMS revealed that pharmacological noradrenergic enhancement via NA agonists *increases* the size of MEPs, indicating that enhanced NA has a facilitatory effect on cortical excitability^16–23^. Noradrenergic suppression, on the other hand, has been shown to lead to a decrease in excitability^24^. Such acute modulatory effects were observed after both single-dose and chronic drug administration and thus do not seem to depend on mechanisms specifically linked to long-term drug application (reviewed in^25^). Second, recent work suggests that global changes in arousal elicited by activity changes in neuromodulatory systems are reflected in non-oscillatory, broadband neural activity that can be captured using scalp-electroencephalography (EEG) in humans^26–31^. Such a broadband component reflects the decay of power with increasing frequency and is known as the 1/f slope of the EEG power spectrum, or “spectral slope”. A flatter slope, reflecting a power shift from low-frequency to high-frequency power, has been associated with increased neuronal activation^26,32,33^. In line with the proposition that these excitability transitions reflect changes in global brain states, the spectral slope could successfully discriminate wakefulness from different sleep stages and anesthesia^27^. Further, it scaled with pupil-linked arousal measures during quiet wake in humans^29^ and has thus been proposed to serve as an electrophysiological marker of arousal-dependent fluctuations in the human brain^27^. Using computational modeling, Gao et al.^26^ proposed that changes in the spectral slope might depend on synaptic excitation (E) and inhibition (I) balances of neural circuits, with higher E/I relating to increased *intrinsic* excitability (i.e., independent of external stimulation) related to a flatter slope^26^.

Theories of LC functioning suggest an interplay of tonic activation patterns, reflecting the brain’s global arousal state, and phasic responses related to task engagement and performance^2,34^. The oddball paradigm, an attention task where participants attend and respond to targets while ignoring frequently presented standard stimuli, has been widely used to probe this interaction. Research in monkeys demonstrates that relatively high tonic activity is associated with smaller phasic responses and lower task performance^35–37^. Both pupil dilation and the P300, a heavily researched event related potential (ERP) component peaking 250-500ms after presentation of task-relevant stimuli such as targets of an oddball task^38^, have been suggested to reflect psychophysiological correlates of the phasic LC response^34,39^. In humans, a relation between pupil size at baseline, suggested to reflect tonic LC activity levels, and pupil dilation responses to target tones, the P300, and task performance has been reported^39–41^. Along similar lines, we could recently show that by combining pupil-BF with an auditory oddball task, self-regulation of baseline pupil size prior to target presentation systematically modulated both pupil dilation responses to target tones and task performance^14^. Whether such self-regulation further influences the P300 has yet to be determined.

Here, we aimed to further spotlight the effects of the recently developed pupil-BF approach on different markers of cortical excitability and arousal in a series of experiments using a multimodal approach (Figure 1a-e). Combining pupil-BF with single-pulse TMS, we investigated whether pupil self-regulation through pupil-BF modulates cortical excitability. Specifically, in line with previous NA drug administration studies^25^, we hypothesized to find increased acute cortical excitability reflected in larger MEPs when participants upregulate pupil size as compared to downregulate pupil size (Experiment 1, Figure 1c). Combining pupil-BF with EEG, we expected that the spectral slope, previously suggested to reflect a global state of cortical arousal and *intrinsic* excitability, is flatter during pupil size upregulation (marking a higher arousal state) as compared to downregulation (Experiment 2, Figure 1d). For both experiments, we investigated whether changes in cortical measures of excitability and arousal are correlated with changes in cardiovascular arousal. Further tapping into proposed theories of LC functioning and the interplay between tonic and phasic LC activity, we examined whether pupil self-regulation during a classical two-tone oddball task modulates (1) the spectral slope reflecting global arousal and (2) the P300 response, an electrophysiological marker of phasic LC responses (Experiment 3, Figure 1e). Specifically, we hypothesized to find the spectral slope not only to be modulated prior to tone onset (as in Experiment 2) but also to change in response to targets, indicating a global increase in arousal when processing task-relevant stimuli. Further, we expected that our previous findings of larger pupil dilation responses to targets and increased task performance during pupil downregulation as compared to upregulation^14^ are further reflected in larger P300 amplitudes in response to targets.

**Figure 1.**
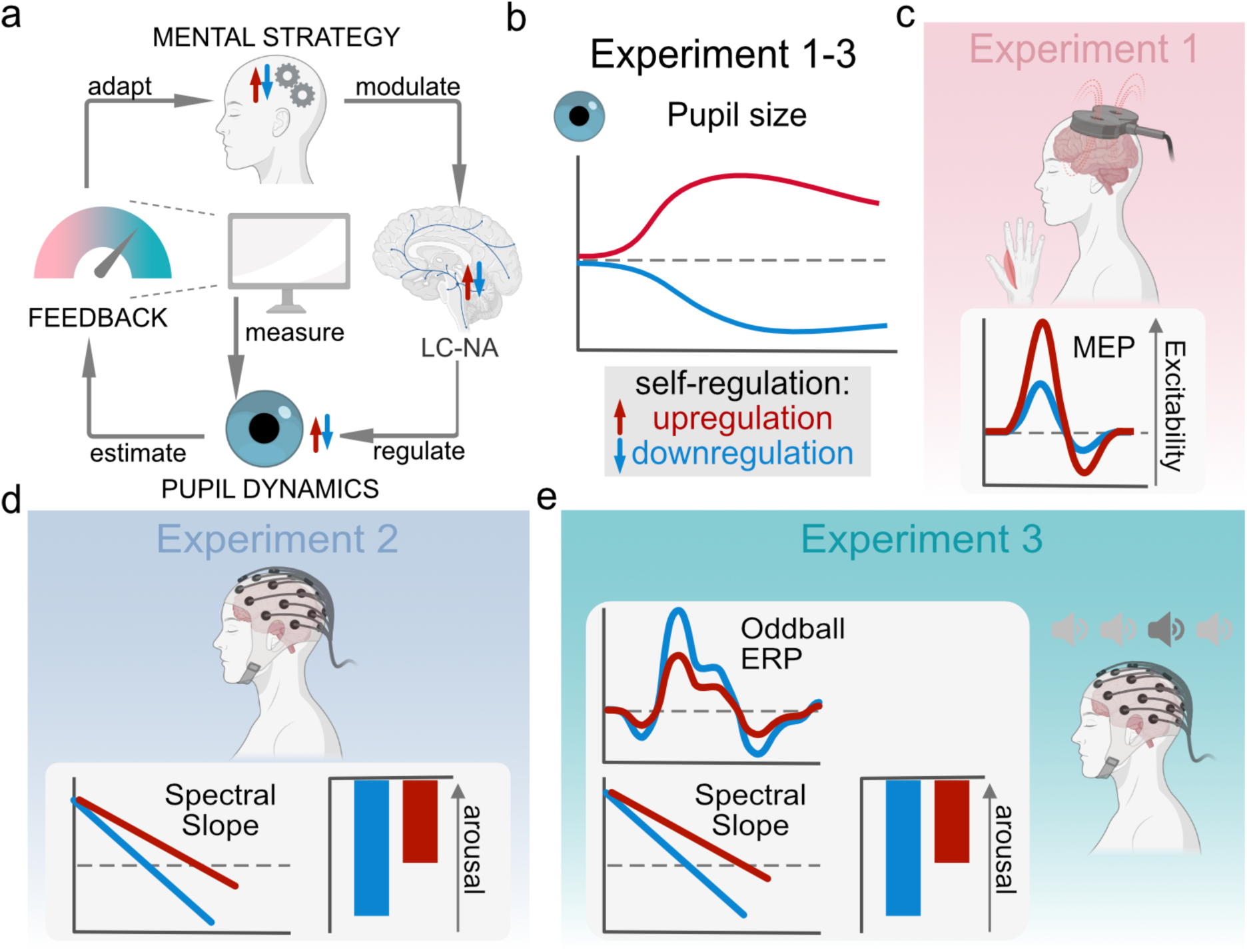
Pupil-based biofeedback (Pupil-BF) approach and its implementation in Experiment 1-3. (a) Pupil-BF approach where participants apply mental strategies to increase (i.e., upregulate) and decrease (i.e., downregulate their pupil size) that have been shown to modulate neuromodulatory systems mediating the brain’s arousal levels such as the LC-NA system. Pupil size is measured by an eye tracker and visually fed back to the participant on a screen. (b) Pupil self-regulation acquired through pupil-BF training in Experiment 1-3. All participants were trained with pupil-based biofeedback in three training sessions on three separate days to learn to volitionally increase (i.e., upregulate) and decrease (i.e., downregulate) their own pupil size. Schematic colored traces indicate pupil size during upregulation (red) and downregulation (blue). (c) Pupil-BF combined with single-pulse TMS to examine how pupil self-regulation influences cortical excitability measured by motor evoked potentials (MEP). We expected increase in cortical excitability, i.e., larger MEP, during upregulation (red) than during downregulation (blue). (d) Pupil-BF combined with EEG to examine how pupil self-regulation influences the spectral slope, a cortical marker of arousal. We expected that the spectral slope, a global marker of cortical arousal is significantly flatter during pupil size upregulation (red) (i.e., indicating higher arousal) as compared to downregulation (blue). The bar graph on the right displays the estimated spectral exponent. (e) Pupil-BF combined with EEG during a simultaneous auditory oddball task to examine how pupil self-regulation influences the spectral slope in response to target tones and the P300 ERP component, an electrophysiological marker of phasic LC activity. Similar to Experiment 2, we hypothesized the slope to be steeper during pupil size downregulation (blue) than upregulation trials (red) and further investigated whether the slope changes in response to target tones. Furthermore, we expected that the P300 would be larger when participants downregulate (blue) as compared to upregulate pupil size (red) prior to target tones. Please note that sound icons in (e) represent standard (light grey) and target tones (dark grey) played during the oddball task. Created with BioRender.com.

## Results

### Experiment 1. Pupil-BF modulates cortical excitability

In Experiment 1, we investigated whether pupil self-regulation through pupil-BF (Figure 1ab) modulates physiological parameters of cortical excitability and how these parameters are linked to cardiovascular markers of arousal in healthy volunteers (Figure 1c). We used a multimodal approach combining pupil-BF^14^, TMS, and electrocardiography (ECG).

We re-recruited 15 participants previously trained with pupil-BF^14^ who underwent an experimental session comprised of 3 conditions (pupil size upregulation, downregulation and resting control; the condition order was counterbalanced across participants). During self-regulation or resting control, cortical excitability was measured with motor evoked potentials (MEPs) elicited by single-pulse TMS. The session consisted of 20 upregulation, 20 downregulation, and 20 control trials with participants self-regulating their pupil size for 15 s (Figure 2). Two TMS pulses were delivered per trial, resulting in 40 MEPs per condition. Additionally, ECG was recorded throughout the experiment to estimate heart rate and HRV in the different conditions.

**Figure 2.**
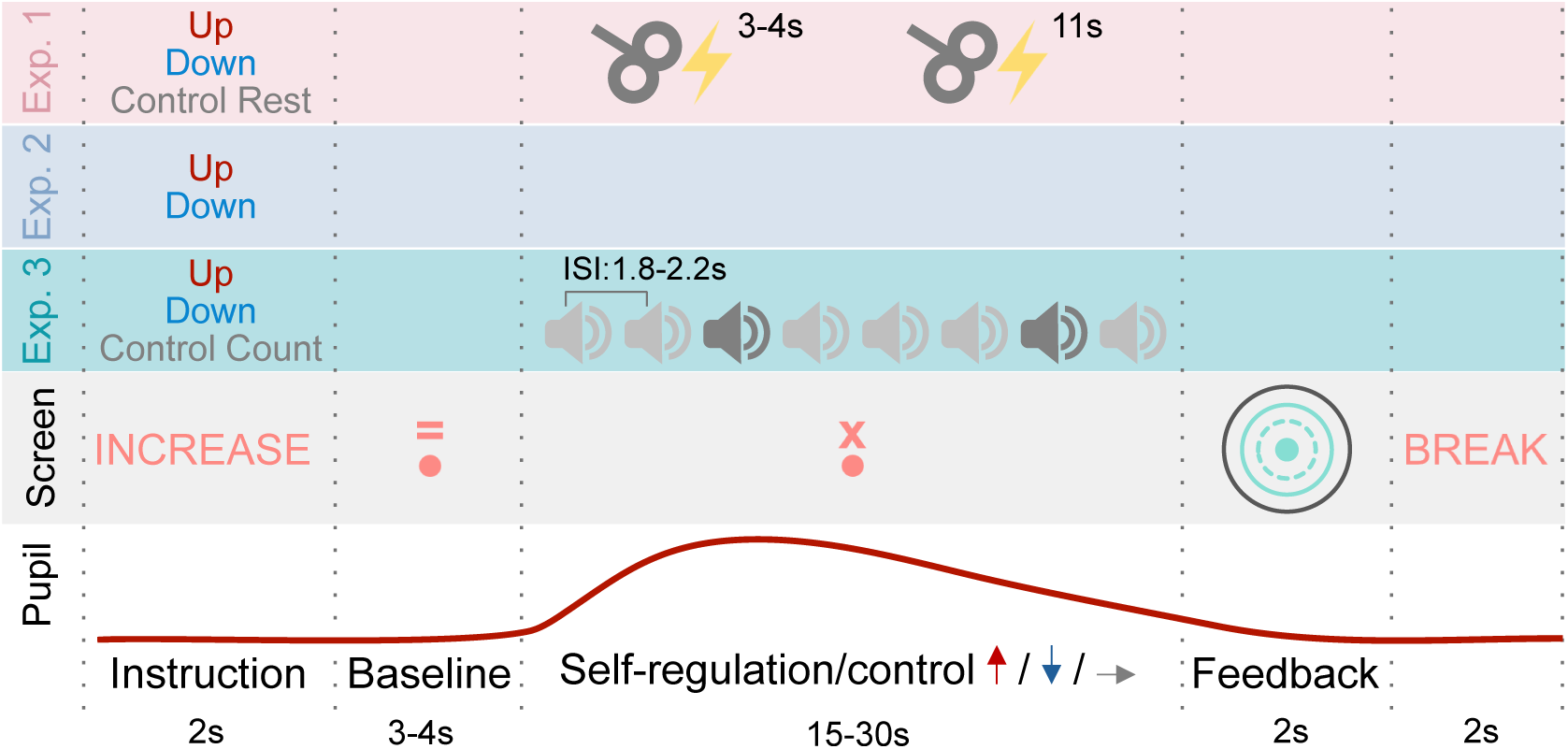
Trial Routine during experiment 1-3. Pupil upregulation, downregulation or a control condition (Experiment 1 and 3) was combined with TMS single pulses (upper panel; Experiment 1), EEG recordings at rest (2^nd^ panel from the top; Experiment 2), or EEG recording during and auditory oddball task (middle panel, Experiment 3). First, participants receive instructions on whether they perform pupil up-, downregulation or a control task, which is followed by a baseline measurement indicated by an ‘=’ above the fixation dot. Then they apply their mental strategies (or the control task; indicated by an ‘x’ above the fixation dot) while TMS pulses are applied (Experiment 1), EEG is recorded (Experiment 2), or target (dark grey sound icon) or standard tones (light grey sound icon) are presented while EEG is recorded (Experiment 3). After this modulation phase, participants received color-coded post-performance feedback (color coding: green in successful and pink in unsuccessful trials, the dotted line indicated baseline pupil size, the solid line indicated average pupil size during modulation and the black like indicated max/min pupil size during modulation) which was followed by a short break before the next trial began (2^nd^ panel from the bottom). The bottom panel indicates exemplary pupil size as measured during the trials, together with the timing of each trial phase.

#### Successful pupil self-regulation under challenging experimental conditions

Study participants were able to voluntarily up- and downregulate their pupil size despite two TMS pulses being applied during the modulation phase (Figure 3a). Replicating previous results^14^, pupil size (Figure 3b) was significantly larger during up- as compared to downregulation (*F*_(2,28)_=14.52, *p*<.001, *η*_p_^2^=0.5; paired samples t-test: *t*_(14)_=4.19; *p*=.003; *d*=1.08; 95%-CI_d_=[0.43;1.71], MD=.33), and control trials (paired samples t-test: *t*_(14)_=4.09; *p*=.002; *d*=1.06; 95%-CI_d_=[0.41;1.68], MD=.27; corrected for multiple comparisons using sequential Bonferroni correction). No significant differences were found between downregulation and control trials (paired samples t-test: *p=.*19, MD=-.07) which is not unexpected because the control conditions required participants to rest and relax while thinking of nothing in particular.

**Figure 3.**
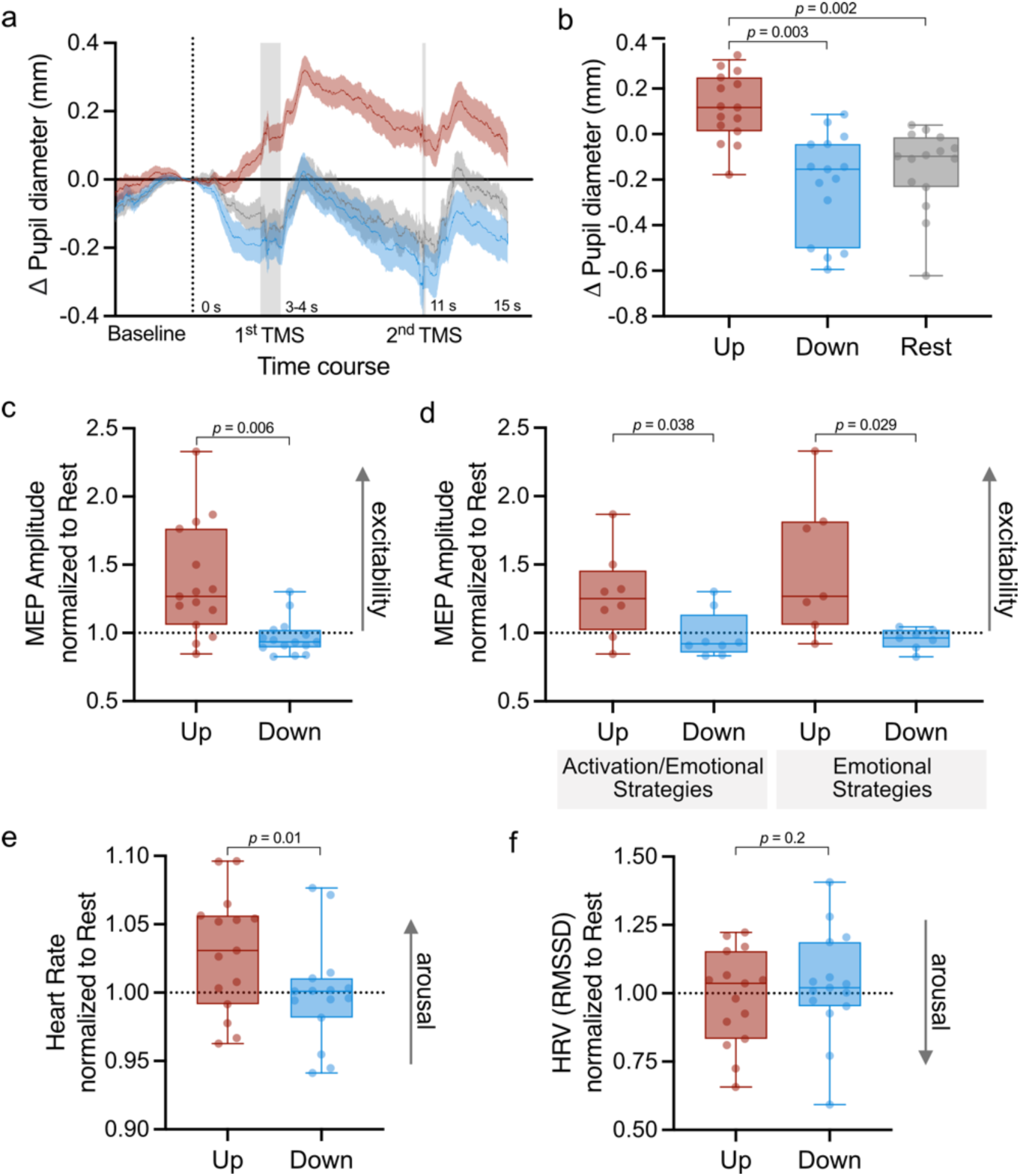
The effects of pupil self-regulation on cortical excitability and cardiovascular arousal. (a) Pupil self-regulation during the application of TMS pulses. A time course of pupil size changes during upregulation (red), downregulation (blue), and rest (control, grey) trials. Shaded grey areas depict the timing of TMS pulses, where the 1^st^ pulse was applied between 3-4s and 2^nd^ pulse 11s after the modulation onset. Note that for the participant the timing of the 2^nd^ pulse was subjectively variable in relation to the 1^st^ pulse. Each trial started with a baseline pupil measurement (until the dotted line) and was followed by a 15s modulation (or rest phase) where 2 TMS pulses were applied to measure the momentary cortical excitability. (b) Pupil size changes during modulation averaged across 150ms before each TMS pulse (baseline-corrected to 1s baseline prior to modulation start) in the upregulation, downregulation, and control rest conditions. During upregulation trials, baseline-corrected pupil size changes were significantly larger as compared to downregulation and control trials (post-hoc paired samples t-tests: up versus down: t(14)=4.19, p=.003; up versus control t(14)=4.09, p=.002; corrected for multiple comparisons using sequential Bonferroni correction). Pupil size changes during downregulation were not significantly lower as compared to control trials. (c) Motor evoked potential (MEP) amplitude measured during up- and downregulation, respectively, displayed normalized to the resting control condition. Pupil size upregulation led to significantly larger MEPs as compared to downregulation (t(14)=10.35; p=.006). (d) Difference of MEPs between up- and downregulation conditions for participants applying activation combined with emotion-related strategies versus purely emotion-related strategies. For both strategy types, MEPs during pupil size upregulation were larger as compared to during pupil size downregulation (activation: t(7)=3.02; p=.038; emotion: t(6)=2.0; p=.029; p-values adjusted with sequential Bonferroni correction). Heart rate (e) and heart rate variability (f) averaged for up- and downregulation trials across all participants, normalized to the resting control condition. Heart rate variability (HRV) was estimated as the root mean square of successive differences (RMSSD). Self-regulation of pupil size systematically modulated heart rate leading to significantly higher heart rates during upregulation as compared to downregulation trials (t(14)=2.9; p=.01). RMSSD, in contrast, was not significantly modulated by pupil self-regulation during simultaneous TMS pulse application. Shaded areas indicate the standard error of the mean (s.e.m). Boxplots indicate median (center line), 25th and 75th percentiles (box), and maximum and minimum values (whiskers). Dots indicate individual participants.

#### Pupil self-regulation modulates cortical excitability

We found that MEP amplitude measured during the modulation phase, normalized to the resting control condition, was higher in the upregulation than in downregulation condition (MD=.36, i.e., 36%; *t*_(1,14)_=10.35, *p*=.006, *η*_p_^2^=.43; Figure 3c). To investigate whether this result was driven by background electromyography (EMG) activity (bgEMG), we added bgEMG activity as a covariate to the statistical model. The effect of self-regulation remained highly significant (*F*_(1,13)_=8.14, *p*=.01, *η*_p_^2^=.39), indicating increased cortical excitability when participants applied strategies to upregulate their pupil size independent of background EMG activity. An additional control analysis using Bayesian repeated measures (rm) analyses of variance (ANOVA) with the factor condition (upregulation, downregulation and control) further supported that background EMG activity did indeed not differ between conditions (BF_01_=4.83, i.e., moderate evidence for the H0, see Supplementary Figure 1).

Since mental strategies to upregulate pupil size could stem from two different categories (i.e., emotional versus physical activation-related strategies), we split the data set into groups using (i) purely emotional strategies and (ii) a mix of physical activation and emotional strategies. This additional step was taken to rule out that effects on cortical excitability were not purely driven by imagining physical activity, as part of activation strategies, which has been shown to activate the primary motor cortex^42,43^. We found that in both groups (i.e., physical activation and emotion-related strategies), upregulation of pupil size led to a significant increase in MEP amplitude in comparison to downregulation (both normalized to control; Figure 3d; physical activation: two-sided paired samples t-test: *t*_(7)_=3.02, *p*=.038, *d*=1.1, 95%-CI_d_=[0.16;1.93]), MD=.29, 29%; emotion-related: two-sided paired samples t-test: *t*_(6)_=2.9, *p*=.029, *d*=1.1, 95%-CI_d_=[0.10;2.01]), MD=.53, 53%; corrected for multiple comparisons using sequential Bonferroni correction). Pupil size and MEP changes were largely unrelated (Pearson’s correlation: *p*=.46; Supplementary Figure 2a).

#### Pupil self-regulation during TMS significantly modulates heart rate but not heart rate variability

We tested the influence of self-regulating pupil size on cardiovascular parameters using ECG signals during pupil upregulation, downregulation and control conditions. We observed higher heart rate during upregulation than downregulation trials (both normalized to the rest condition; two-sided paired samples t-test: *t*_(14)_=2.9, *p*=.01, *d*=.74, 95%-CI_d_=[0.15;1.3]), MD=.03, Figure 3e). To assess HRV, we calculated the root mean square of successive differences (RMSSD). The RMSSD was generally higher during downregulation than during upregulation, however there was no significant effect of pupil self-regulation on RMSSD (both normalized to the rest condition; two-sided paired samples t-test: *t_(_*_14)_=-1.3, *p*=0.2, *d*=-.34, 95%-CI_d_=[-0.86;0.18], MD=-.05, Figure 3f). MEP and changes in heart rate and HRV were largely unrelated (Pearson’s correlation: *p*=.92 and *p*=.3, respectively; Supplementary Figure 2bc).

### Experiment 2. Pupil-BF modulates an electrophysiological marker of cortical arousal

To test the influence of pupil-BF on cortical and cardiovascular arousal markers, twenty-five participants were recruited via online advertisement or re-recruited from a pool of previously trained participants^14^. All participants underwent three days of pupil-BF training of which 23 participants completed all three days. In case of re-recruitment of participants, they received one session of re-training that was similar to training day 3 of newly recruited participants. Before the first experimental session, all participants received standardized instructions on the mental strategies for self-regulating pupil size, which were derived from previous studies (for details, see Meissner et al.^14^). Each session was composed of 30 upregulation and 30 downregulation trials, with participants aiming to modulate their pupil diameter for 15s (day 1 and day 2) or 30s (day 3) while receiving isoluminant visual real-time (day 1 and day 2) and post-trial performance feedback (all days).

#### Pupil self-regulation through pupil-BF training

Here, we focus on pupil self-regulation data of day 3 not reported before. To be consistent throughout all experiments, we report data of the first 15s of modulation (for data analysis of the complete time window of 30s, see Supplementary Figure 3). Pupil self-regulation was quantified by calculating a baseline-corrected upregulation and downregulation score (i.e., average pupil size change during upregulation and downregulation, respectively, as compared to baseline, Figure 4a) and a pupil modulation index (upregulation-downregulation, i.e., the difference between pupil diameter changes in the two conditions, Figure 4b). The pupil modulation index was significantly larger than 0 (two-sided one sample t-test: *t*_(22)_=4.85, *p*<.001, *d*=1.01; 95%-CI_d_=[0.50;1.51]), indicating participants’ ability to voluntarily modulate pupil size at the third day of training. This successful modulation on day 3 was mainly driven by a strong downregulation (two-sided one sample t-test downregulation: *t*_(22)_=-6.01, *p*<.001, *d*=-1.25; 95%-CI_d_=[-1.80;-0.70]; upregulation: *p*=.15; corrected for multiple comparisons using sequential Bonferroni correction; Figure 4a).

**Figure 4.**
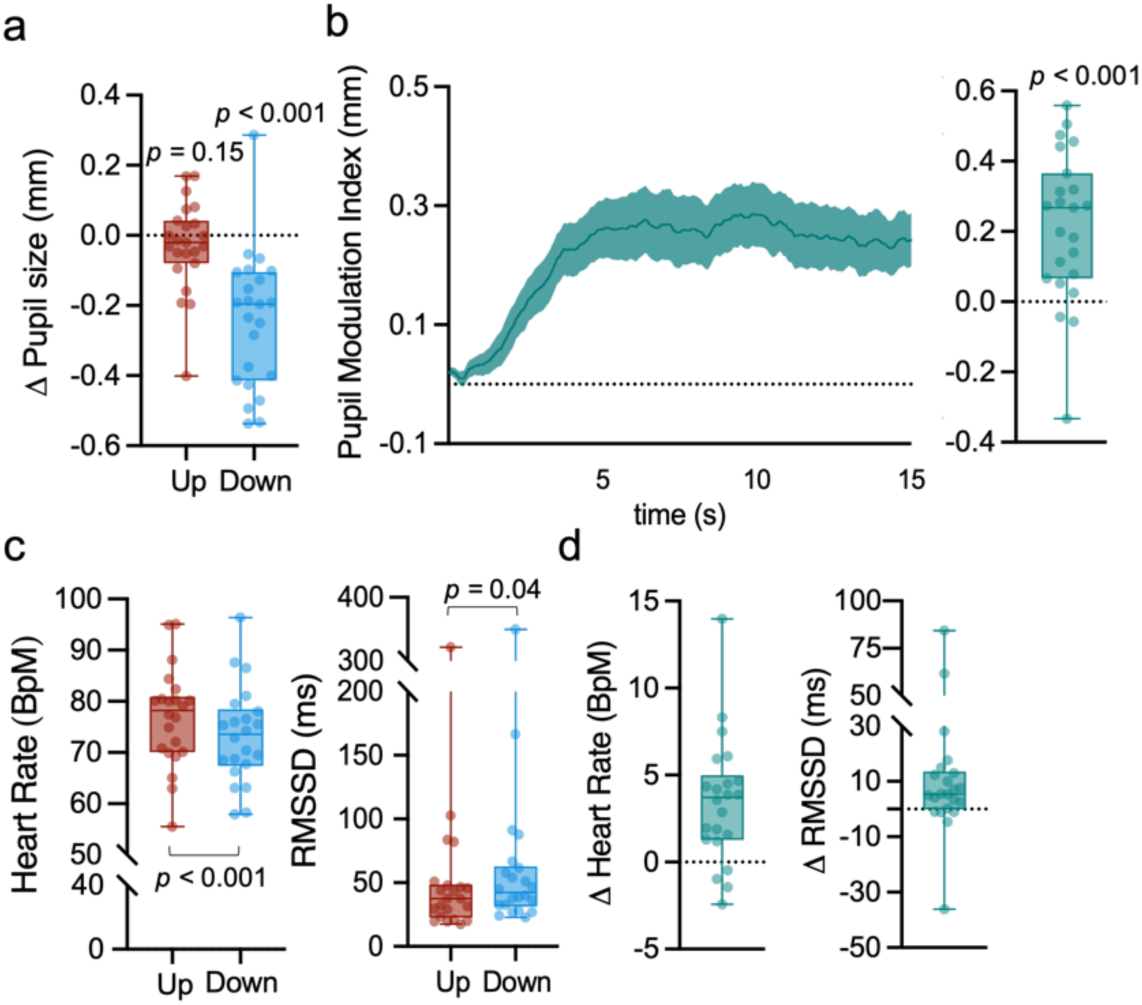
Effects of pupil self-regulation on pupil size and cardiovascular arousal at the end of pupil-BF training. (a) Pupil upregulation and downregulation, respectively, averaged across the 15s modulation phase. We found a significant difference between pupil size downregulation as compared to upregulation (paired samples t-test: t(22)=-6.01; p<.001). (b) Time series of the pupil modulation index reflecting the difference between the average pupil size during the two conditions (Up–Down; left panel). The pupil modulation index averaged across the 15s modulation phase where larger values indicate better ability to self-regulate pupil size. (c) Heart rate and heart rate variability (HRV) averaged for up- and downregulation trials across all participants. HRV was estimated as the root mean square of successive differences (RMSSD). Self-regulation of pupil size systematically modulated heart rate and RMSSD leading to significantly higher heart rates (paired samples t-test: t(21)=-4.45; p<.001) and lower RMSSD (paired samples t-test: t(21)=-2.17; p=.04) during upregulation as compared to downregulation trials. Shaded areas indicate the standard error of the mean (s.e.m). Boxplots indicate median (center line), 25th and 75th percentiles (box), and maximum and minimum values (whiskers). Dots indicate individual participants.

#### Pupil self-regulation modulates the spectral slope, a marker of cortical arousal

Next, we investigated whether pupil self-regulation modulates the spectral slope, the aperiodic component of the EEG power spectrum (Figure 5a) which has recently been suggested to represent an electrophysiological marker of arousal in humans^27,29^. As hypothesized, the spectral slope estimated for 30-43Hz during downregulation was significantly steeper than during upregulation (two-sided paired samples t-test: *t_(22)_*=3.37; *p=*.003; *d=*.70; 95%-CI_d_=[0.24;1.16]; Figure 5b); for similar trend-level results of the spectral slope estimated for 2-40Hz, see Supplementary Figure 4ab), suggesting a lower state of cortical arousal during downregulation as compared to upregulation of pupil size. Importantly, this difference was present during self-regulation but not during baseline (rmANOVA: time bin*condition interaction: *F*_(4.01,88.26)_=2.52, *p=*.047; *η*_p_^2^=.10; 95%-CIη_p_^2^=[0.00;0.20]; post-hoc paired samples t-tests: Baseline: *p=*.97; 0-3s: *t_(22)_*=3.33; *p=*.018; *d=*.70; 95%-CI_d_=[0.23;1.15]; 3-6s: *t_(22)_*=2.39; *p=*.05; *d=*.50; 95%-CI_d_ =[0.06;0.93]; 6-9s: *t_(22)_*=2.61; *p=*.06; *d=*.54; 95%-CI_d_ =[0.10;0.98]; 9-12s: *t_(22)_*=2.95; *p=*.035; *d=*.62; 95%-CI_d_=[0.16;1.06]; 12-15s: *t_(22)_*=2.43; *p=*.07; *d=*.51; 95%-CI_d_ =[0.07;0.94]; corrected for multiple comparisons using sequential Bonferroni correction; Figure 5c), indicating a strong modulatory effect of pupil self-regulation on global cortical arousal measured with EEG. Including all participants, pupil size and spectral slope changes were largely unrelated (Pearson’s correlation: *p=*.70; Supplementary Figure 4c). However, this was mainly driven by an outlier with a very low pupil modulation index. Removing this outlier, we found a significant positive relationship between the pupil modulation index (upregulation-downregulation) and spectral slope changes (upregulation-downregulation; *r*=0.43; *p*=.048; 95%-CI_r_=[0.01;0.72]; Figure 5d), thus better pupil self-regulation was associated with larger differences in spectral slope.

**Figure 5.**
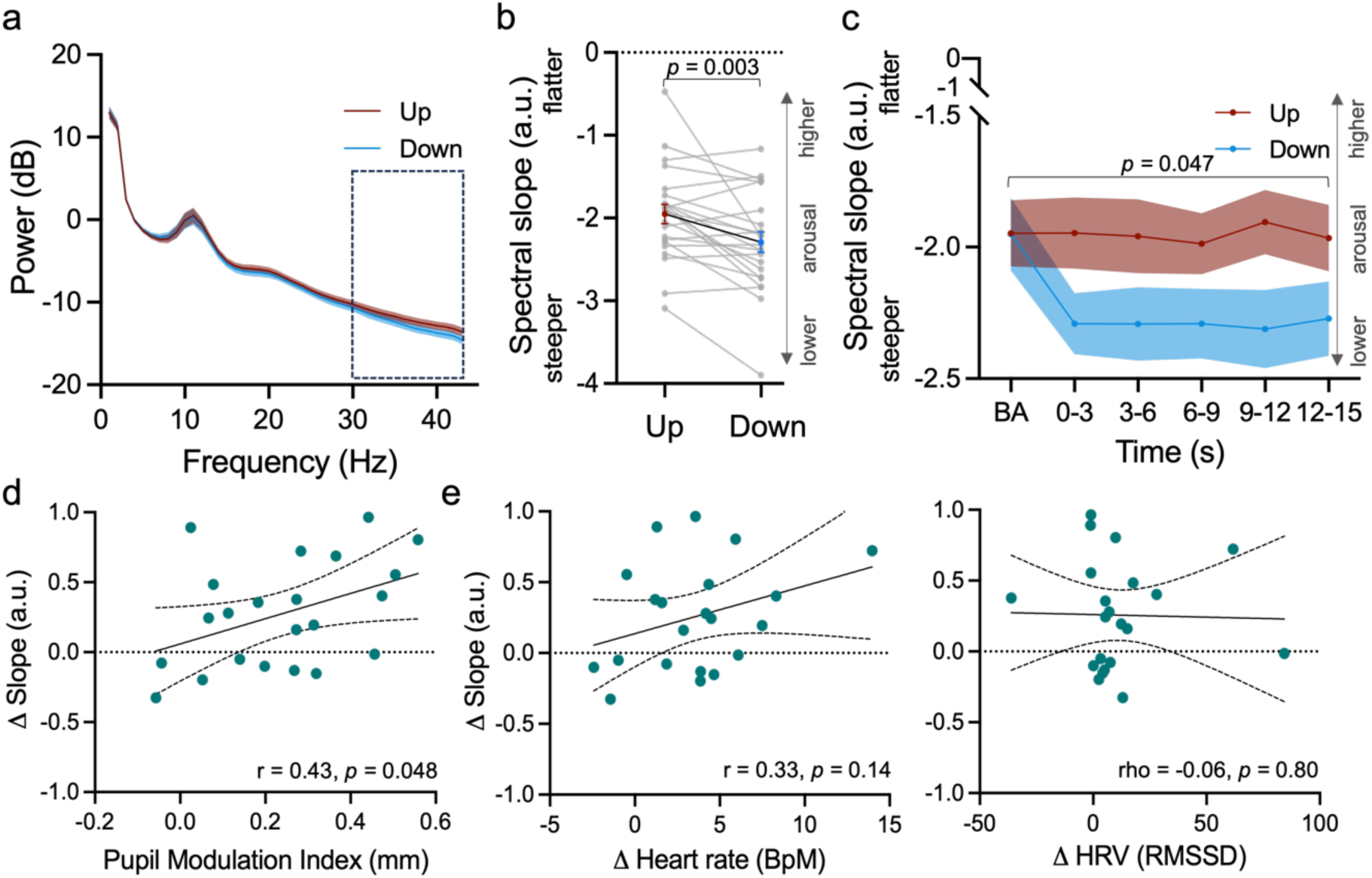
Effects of pupil self-regulation on the spectral slope. (a) Power spectra averaged across all channels for up- and downregulation trials. Dashed box outlines frequency range (i.e., 30-43 Hz) for which slope is estimated and reported throughout the main manuscript. (b) The spectral slope estimated for 30-43 Hz averaged across all channels and time windows of pupil size up- (red) and downregulation (blue), indicating significantly steeper slopes during down- as compared to upregulation (t(22) = 3.37; p = 0.003). (c) The spectral slope estimated for 30-43 Hz averaged across all channels of up- (red) and downregulation (blue) trials for time bins of 3s during baseline (i.e., -3-0s) and the modulation phase (0-3s, 3-6s, 6-9s, 9-12s, 12-15s). The spectral slope was significantly steeper during downregulation during most of the modulation time windows, however, not during baseline recordings (time bin*self-regulation interaction: F(4.01,88.26)=2.52, p=.047). (d) Pearson correlation coefficient between pupil modulation indices (i.e., the difference between pupil size changes in up-versus downregulation trials, Up-Down) and differences in spectral slope between up- and downregulation trials averaged across the 15s modulation phase, indicating a significant linear relationship between pupillary and cortical markers of arousal. (e) Non-significant Pearson (left panel) and Spearman rho correlation coefficients (right panel) between differences in spectral slope (upregulation-downregulation) and differences in heart rate (left panel; upregulation-downregulation) and RMSSD (right panel; downregulation-upregulation). Shaded areas indicate s.e.m. Dots indicate individual participants.

#### Pupil self-regulation modulates cardiovascular arousal markers

To examine the influence of pupil self-regulation on cardiovascular parameters, we analyzed ECG data during pupil self-regulation on day 3 (*n*=22, see Methods for exclusion of 1 participant). Similar to previous results, we observed significanlty higher heart rate during upregulation versus downregulation trials (paired samples t-test: *t*_(21)_=4.45; *p*<.001; *d*=0.95; 95%-CI_d_=[0.44;1.45], Figure 4cd). To assess HRV during pupil self-regulation, we calculated the RMSSD. We found a significant effect of pupil self-regulation on RMSSD with larger RMSSD values during downregulation as compared to upregulation (paired samples t-test: *t*_(21)_=-2.17; *p*=.04; *d*=-0.46; 95%-CI_d_=[-0.90;-0.02], Figure 4cd), suggesting higher parasympathetic activity during downregulation as compared to upregulation of pupil size.

#### There is no significant linear relationship between cortical and cardiovascular markers of arousal

Finally, we investigated whether changes in cortical arousal (i.e., the spectral slope, upregulation-downregulation) are directly linked with changes in cardiovascular arousal (i.e., heart rate (upregulation-downregulation) and RMSSD (downregulation-upregulation)). However, neither heart rate (all *p>*.14; Figure 5e, left panel) nor RMSSD changes (all *p>*.44; Figure 5e, right panel) showed a significant linear relationship with spectral slope changes, independent of the removal of the outlier (see *pupil self-regulation modulates the spectral slope;* Supplementary Figure 4d).

### Experiment 3. Pupil-BF modulates electrophysiological markers of LC-NA activity and cortical arousal during the auditory oddball task

In Experiment 3, we investigated whether pupil self-regulation modulates EEG markers of arousal when combined with a two-tone auditory oddball task. We assessed the spectral slope, a marker previously linked to global arousal state^27^, and the P300, an electrophysiological marker previously linked to phasic LC activity^34,39^. The data on pupillary responses and behavioral performance have been previously published^14^, thus, we focus on EEG dynamics here. 22 trained participants performed the oddball task while (i) upregulating pupil size (ii) downregulating pupil size or (iii) executing a cognitive control task of silently counting backwards in steps of seven. The control task was included to control for cognitive effort effects that may arise from simultaneously executing pupil self-regulation and the oddball task. In general, participants (n = 20 for pupillary analyses) were able to successfully self-regulate pupil size in this dual task setup, which has been previously published^14^. Of these, 19 participants were included for EEG analyses (see Methods section).

#### Pupil self-regulation modulates the spectral slope during the oddball task

First, we investigated whether pupil self-regulation modulates the spectral slope during the auditory oddball task. As expected, the spectral slope was significantly steeper during downregulation as compared to upregulation and control trials, independent of the sound category (rmANOVA; main effect of condition: *F*_(2,36)_=21.08; *p*<.001; η_p_^2^=0.54; 95%-CIη_p_^2^=[0.32;0.65]; post-hoc paired samples t-tests, corrected for multiple comparisons using sequential Bonferroni correction: upregulation vs. downregulation: *t*_(19)_=-6.42; *p*<.001; *d*=-1.43; 95%-CI_d_=[-2.06;-0.80]); upregulation vs. control: *t*_(18)_=-.37; *p*=.74; *d*= -0.08; 95%-CI_d_=[-0.53;0.37], Figure 6a); downregulation vs. control: *t*_(18)_=4.55; *p*<.001; *d*=1.04; 95%-CI_d_=[0.47;1.60]). Furthermore, this analysis revealed a significant time bin*sound interaction (*F*_(5,90)_=3.30; *p*=.009; η_p_^2^=0.16; 95%-CIη_p_^2^=[0.02;0.23]) with significantly flatter slopes following target (i.e., 500-1000ms; *t_(_*_18)_=2.96; *p=*.04; *d*=0.68; 95%-CI_d_=[0.17;1.17]; corrected for multiple comparisons; Figure 6ab) but not prior to target as compared to standard sounds (*p*=.53). This effect was, however, independent of the self-regulation condition (i.e., no significant interaction effect with the factor condition; all *p>.*20).

**Figure 6.**
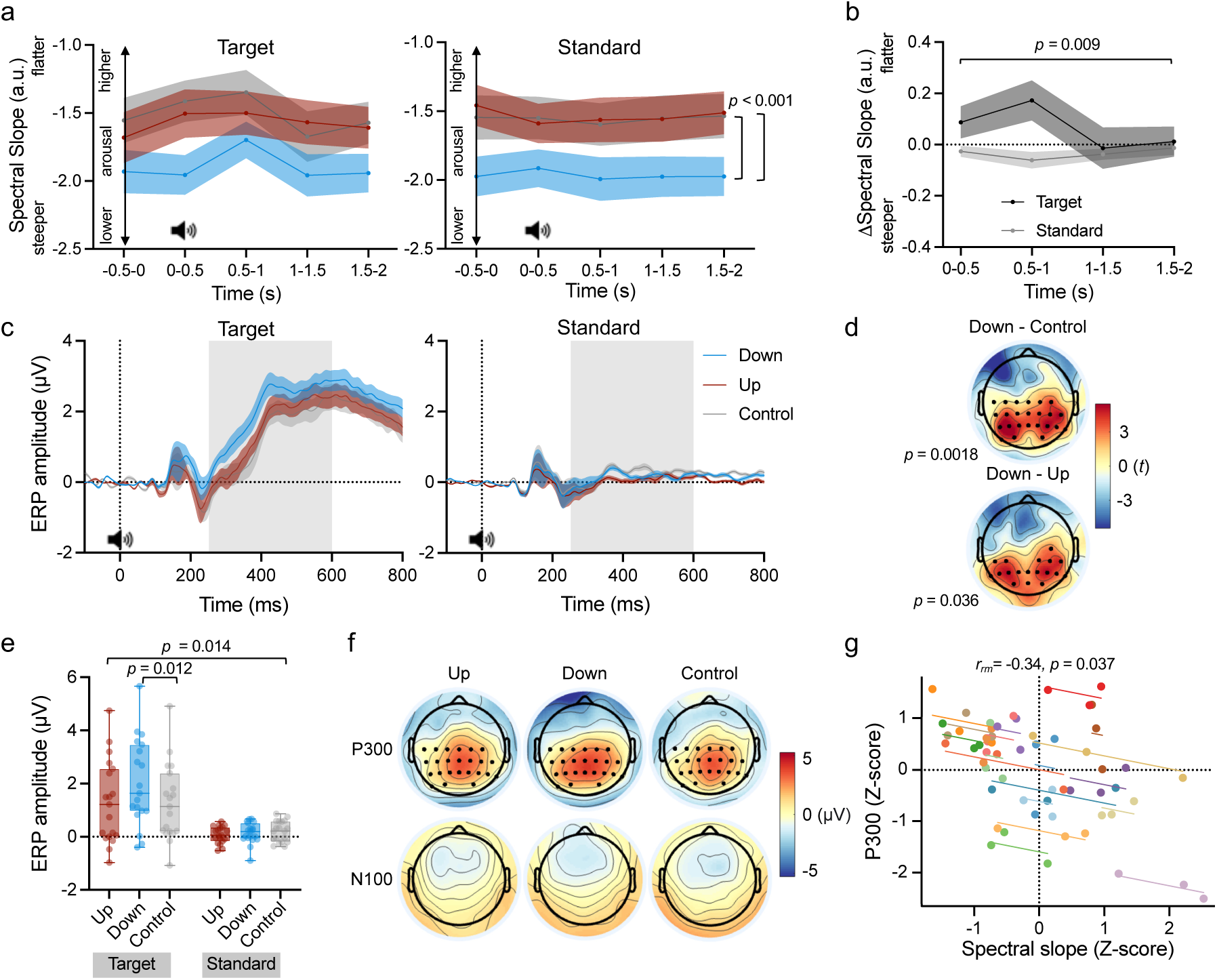
Effects of pupil self-regulation on the spectral slope, P300, and N100 during the oddball task. (a) The spectral slope estimated for 30-43 Hz averaged across all channels of up- (red), downregulation (blue), and control trials for time bins of 500s prior to (−500-0ms) and after target and standard tone onset (i.e., 0-500ms, 500-1000ms, 1000-1500ms, 1500-2000ms). The spectral slope was significantly steeper during downregulation as compared to upregulation and control trials (main effect of condition: F(2,36)=21.08, p<.001) independent of whether targets or standards were played. (b) Differences in the spectral slope between the time windows following tone onset and the time window prior to tone onset (i.e., post-pre). Please note that these differences depicted here are averaged across all three conditions and illustrates that the spectral slope was flattened in response to targets as compared to standard sounds (time bin*sound category interaction: F(5,90)=3.30; p=.009), an effect that is independent of the self-regulation condition. Note that statistics are reported for the full model will all time bins and conditions, however, here we show simplified baseline-corrected differences averaged across conditions for illustrative purposes. (c) ERP time series averaged across the most prominent channels (C3, CP5, CP1, Pz, P3, P4, P8, CP6, CP2, C4, C1, C5, CP3, P1, P5, PO7, PO3, PO4, P6, P2, CPz, CP4, C2, Cz). The grey box indicates the time window (252-600ms) for which the P300 was compared between the modulation conditions. (d) Resulting t-values of the cluster-based permutation t-tests show a difference in the P300 between downregulation and control (t=2202.5, p=.0018; top) and downregulation and upregulation (t=825.7, p=.036; bottom) across central and posterior channels (most prominent channels are marked with a black dot). (e) There was a significant sound*condition interaction on the P300 (F(2,36)=4.82; p=.014, ηp^2^=.21, 95%-CIηp^2^=[.01-.40]), which was driven by a significant difference between the downregulation and cognitive control condition for targets (t(18)=3.52; p=.012, d=.81; 95%-CId=[0.28;1.32]). There were no significant differences for target sounds between downregulation and upregulation (t(18)=-2.70; p=.075) or upregulation and control (t(18)=0.01; p=1). The difference in P300 was specific to targets and did not differ significantly between any of the modulation conditions for standard sounds (all ps>.34). (f) Scalp ERPs averaged over the time window of interest for the P300 (252-600ms; top) and N100 (50-200ms; bottom). Cluster-based permutation t-tests did not show any significant differences between the conditions or standard and target sounds for the N100. (g) A repeated measures correlation shows that baseline spectral slope (−500-0ms) negatively correlates with the P300 amplitude (i.e. a flatter slope (higher arousal) relates to lower P300 amplitudes; rrm=-.34, p=.037). Shaded areas represent s.e.m. Boxplots indicate median (center line), 25th and 75th percentiles (box), and maximum and minimum values (whiskers). Dots and colors indicate individual participants. Sound icon indicates the presentation of a target or standard sound.

#### Pupil self-regulation modulates the P300

Next, we assessed whether pupil-self regulation modulated the P300 in response to target tones during the oddball task. We found that the P300 differed significantly between the downregulation and cognitive control condition (cluster-based permutation t-test on ERPs; 252 – 600ms after target sound onset: *t*=2202.5, *p*=.002, Figure 6c-e). This difference was characterized by a higher amplitude in the downregulation as compared to the control condition across a broad centroparietal cluster and was most prominent from 252 to 560ms after sound onset, corresponding to the expected temporal and topographic properties of the P300. The P300 between the downregulation and upregulation condition also differed significantly (*t*=825.7, *p*=.036). This effect followed a similar pattern, with a higher amplitude in the downregulation than upregulation condition and the most pronounced difference in a centroparietal cluster from 272 – 368ms after tone onset. We did not observe a significant difference in the P300 between the upregulation and control condition. The specificity of the pupil self-regulation modulation effect on the target-evoked P300 was further confirmed by comparing P300 responses to standard and target sounds. In general, the mean amplitude across the centroparietal cluster from 252 to 600ms was significantly higher for target sounds than standard sounds (rmANOVA, main effect of sound: *F*_(1,18)_=26.70, *p*<.001, η_p_^2^=.60, 95%-CIη_p_^2^=[0.24;0.75], Figure 6e). Importantly, there was a significant sound*condition interaction (*F*_(2,36)_=4.82; *p*=.014, η_p_^2^=.21, 95%-CIη_p_^2^=[0.01;0.40]), which was driven by a difference in mean amplitude for target, but not standard sounds, in the downregulation compared to the control condition (post-hoc t-tests: target up vs. target control (*t*_(18)_=0.01; *p*=1); target down vs. target control (*t*_(18)_=3.52; *p*=.012, *d*=.81; 95%-CI_d_=[0.28;1.32]); target up vs. target down (*t*_(18)_=-2.70; *p*=.075); standard up vs. standard control (*t*_(18)_=-1.82, *p*=.34); standard down vs. standard control (t_(18)_=-.78, p=1); standard up vs. standard down: (*t*_(18)_=-0.94, *p*=1); corrected for multiple comparisons using sequential Bonferroni correction). To ensure that the effect of modulation condition was specific to the target-evoked P300, we further tested differences in the N100 between the three conditions but found no significant differences (all *p*>.1; cluster-based permutation t-tests on ERPs from 50 – 200ms after target sound onset, Figure 6f).

#### Cortical arousal before stimulus presentation correlates with P300 responses

Finally, we assessed whether the spectral slope prior to target sound presentation, representing the global cortical arousal state relates to the target-evoked P300 amplitude. A repeated-measures correlation between z-scored spectral slope prior to tone onset (−500-0ms) and z-scored P300 amplitudes in the 252-600ms time window across most prominent channels (Figure 6d) showed a significant negative correlation (*r_rm_*=-.34, *p*=.037, 95%-Ci*r*_rm_=[-0.59;-0.02]; Figure 6g). Thus, a flatter slope (i.e., higher cortical arousal) at baseline relates to larger P300 responses to oddball targets.

## Discussion

In a series of experiments, we investigated whether pupil self-regulation, previously shown to tap into arousal-regulating centers in the brainstem including the LC-NA system^14^, modulates electrophysiological markers of cortical excitability, cortical arousal and phasic LC activity. Combining pupil-BF with (i) single-pulse TMS, (ii) EEG recordings, and (iii) a simultaneous auditory oddball task revealed three main findings: First, pupil self-regulation significantly modulated (i) cortical excitability measured with TMS (Figure 3) and (ii) the slope of the EEG power spectrum (i.e., spectral slope), an electrophysiological marker of cortical arousal in humans, proposed to reflect intrinsic excitability via capturing shifts in E/I ratios^26,27^ (Figure 5). Second, during the oddball task, the spectral slope flattened in response to target tones, indicating the sensitivity of the slope to attention-related processes (Figure 6ab). However, these changes were not significantly modulated by pupil self-regulation. Third, tapping further into postulated LC-NA functioning, pupil self-regulation significantly modulated the P300 response elicited by target tones (Figure 6d-f). This corroborates our previous findings of larger pupil dilation responses during pupil size downregulation of the same dataset^14^. Based on our findings of the modulation of LC activity via pupil self-regulation, it further supports the interpretation that the P300 may serve as an electrophysiological readout of phasic LC activity^34,44,45^.

We demonstrated that pupil self-regulation significantly modulated cortical excitability as indexed by increased MEPs during pupil size upregulation as compared to downregulation, independently of the used mental strategies (see Figure 3cd). Based on our previous fMRI study, we know that pupil size up-versus downregulation significantly increased and decreased LC activity^14^. Our findings of a significant modulation of cortical excitability here therefore corroborate pharmacological studies observing that drugs enhancing noradrenergic concentrations increase cortical excitability (for a review, see^25^), whereas drugs reducing noradrenergic transmission, acutely reduce excitability of the human motor cortex^24^. Consistently, pupil size upregulation led to a flatter slope of the EEG power spectrum as compared to pupil size downregulation, both when estimating the spectral slope in a narrowband (i.e., 30-43Hz^27,28^; Figure 5a-c) and broadband frequency range (i.e., 2-40Hz^29^, trend-level effects; Supplementary Figure 4ab). This finding is quite remarkable, since large modulations of the spectral slope have been previously reported in participants in drastically reduced states of arousal such as when undergoing anesthesia or comparing different sleep stages to awake states^27^. Here, we could show that pupil self-regulation can lead to similar changes in the spectral slope. Importantly, the algorithm used here to parameterize the spectral slope allows to clearly separate changes in spectral exponents from other arousal-related alterations such as oscillatory power changes in different frequency bands^46^. Together, these results suggest that pupil-linked self-regulation previously shown to modulate human LC-NA activity systematically influences both classical measures of cortical excitability (i.e., MEPs following single-pulse TMS application)^47^ as well as a more recently suggested marker of global arousal reflecting *intrinsic* cortical excitability (e.g., the spectral slope)^26,27,29,48^, proposed to index the balance between excitatory and inhibitory activity of neural circuits^26^.

The pupil modulation index (i.e., the difference between pupil size upregulation and downregulation) reflecting pupil self-regulation ability was linearly linked to differences in the spectral slope between up- and downregulation trials (Figure 5d), suggesting that increases and decreases in pupil-linked arousal may parallel increases and decreases in E/I ratios in cortical neural circuits, respectively. Our findings are in line with previous results reporting a tight relationship between pupil-based arousal measures and the spectral slope in humans, both during quiet wake^29^ and non-rapid eye movement sleep stages^49^. Thus, the spectral slope and pupil size may reflect partially shared physiological mechanisms. This interpretation aligns with previous evidence that systematically activating the LC-NA system not only causes rapid pupil dilation in rodent models^12,13^ but also that NA-enhancing drugs increase the E/I ratios in the cortex, as demonstrated by computational modeling in humans^50^. Here we show that both is modulated via pupil self-regulation through pupil-BF previously related to activity changes in the LC-NA system^14^.

Differences in MEP amplitudes between pupil size up- and downregulation conditions, on the other hand, did not significantly scale with the ability to self-regulate pupil size (i.e., the pupil modulation index; Supplementary Figure 2a). Whether the lack of a significant relationship between changes in cortical excitability measured with TMS and pupil-linked arousal self-regulation is due to inherently different physiological mechanisms or whether it is driven by methodological constraints, such as restricting pupil size analyses to time windows prior to TMS pulse application to exclude a confound by phasic pupil dilation responses to the pulse, remains to be determined. Even though there is, to the best of our knowledge, no study linking single-pulse TMS responses to changes in pupil diameter, a recent study comparing TMS-based and EEG-based cortical excitability markers reported no significant linear relationship between changes in MEPs and the slope of the EEG power spectrum^48^. Thus, cortical excitability measured with TMS may reflect different physiological mechanisms as captured by pupil or spectral slope measures. Similarly, even though we observed a significant modulation of heart rate (Experiment 1 and 2; Figure 3e and 4c) and HRV (Experiment 2; Figure 4c) through pupil self-regulation, these changes in cardiovascular arousal were not significantly related to induced changes in neither MEP amplitude (Supplementary Figure 2bc) nor spectral slope; Figure 4d). Especially the lack of a significant link between cardiac measures and spectral slope is in line with a previous study showing little to no association between heart rate and spectral slope dynamics during human sleep^49^, indicating that cardiovascular measures are not necessarily linearly linked to cortical excitability and cortical arousal dynamics.

Replicating the influence of pupil self-regulation on spectral slope estimates, we observed a similar modulation when combining pupil self-regulation through pupil-BF with an auditory oddball task: the spectral slope was significantly steeper during pupil size downregulation as compared to upregulation and counting control trials, independent of the sound category (i.e., target versus standard sounds; Figure 6a-c). This finding extends our previous analysis of the same dataset^14^, which revealed significantly larger absolute pupil size during upregulation and control conditions as compared to downregulation, indicating lower states of pupil-linked arousal in the latter condition. Taken together, these results further support the interpretation that pupil-linked and spectral slope-based arousal markers may reflect similar physiological mechanisms. Given that the spectral slope captured volitionally induced global changes in pupil-linked arousal, we further sought to test its sensitivity to attention-related processes during the auditory oddball task and its potential interactions with pupil self-regulation^14^. Research in non-human primates suggests that the allocation of attention leads to a shift in the E/I ratio^51^, which should be reflected in a systematic change in spectral slope. Here, we indeed found a significant modulation of the spectral slope during the auditory oddball task: It was flatter following the presentation of targets (i.e., in a time bin of 500-1000ms after tone onset) as compared to standard sounds, suggesting that this flattening is associated with attentional processes (Figure 6ab). This is well in line with a recent study in humans observing a similar flattening of the spectral slope when allocating selective attention^52^ and suggests that task engagement and resources needed to perform the task, especially in the dual task context here, may lead to stronger excitatory brain activity^51^. Importantly, the stimulus-evoked changes in spectral slope were not significantly modulated by pupil self-regulation and were thus independent of pupil-linked arousal levels. Whether EEG spectra are not sensitive enough to capture changes in attention-related processes as a function of global arousal levels or whether our study powered to detect general differences in global arousal levels and psychophysiological readouts such as pupil dilation responses and the P300 remains to be determined. Notably, a recent study^53^ found an attention-dependent *increase*, as opposed to the reported *decrease,* in the steepness of the spectral slope during an auditory oddball task in humans. Those findings, however, may be driven by how spectral slope was calculated: the spectral slope was estimated for a frequency range between 2 and 25Hz^53^. Here, we targeted the narrowband frequency range of 30 to 43Hz in the context of a dual task. Such narrowband slope estimates have been shown to track behavioral performance on an auditory attention task more consistently than broadband estimates (i.e., 1-45Hz) and may indicate that narrowband activity above 30Hz may be a particularly sensitive marker for task-dependent arousal fluctuations linked to behavioral task performance^28^.

Theories of LC functioning suggest an interplay of tonic LC activation patterns, reflecting the global arousal state of the brain, and phasic LC responses to salient stimuli, associated with a network reset facilitating the refocusing of attention^2,34^. Research in monkeys demonstrated that relatively high tonic activity is associated with smaller phasic responses, whereas when tonic activity is relatively lower at an intermediate level, phasic responses are larger^35–37^. A similar relationship has been observed in human pupil measurements, reporting an inverse or inverted-U relationship between naturally fluctuating baseline pupil size and pupil dilation responses^39,40,54^. Consistently, we have previously shown that pupil size downregulation, associated with relatively lower LC activity, led to larger pupil dilation responses and better task performance than a counting control task and pupil size upregulation^14^. One concern regarding pupil dilation responses is that mechanisms specific to the structure of the eye might limit pupil dilation responses if baseline pupil diameter is already large. Here, we therefore investigated another proposed marker of the phasic LC response^34,44,45^, the P300, and its potential modulation via pupil-BF. The P300, a late ERP component peaking 250-500ms after task-relevant stimuli^38^, is a heavily researched component that has been related to mechanisms of attention and memory operations^45^ and incriminated in psychiatric disorders such as attention deficit hyperactivity disorder^55^. Intriguingly, we could show that the P300 in response to targets (but not standards) was significantly larger during pupil size downregulation than upregulation and counting control trials in a large cluster of electrodes over centroparietal brain regions between 252 and 600ms after target onset (Figure 6cd). In contrast, the N100, included as a control component, was not significantly modulated by pupil self-regulation (Figure 6f). This is in line with previous studies reporting a negative (or inverted-U) relationship between randomly fluctuating baseline pupil size and P300 responses^39,40^ and emphasizes the unique sensitivity of the P300 to volitionally induced changes in global arousal levels. The observation that flatter spectral slopes (higher global arousal) prior to stimulus onset were related to smaller P300 amplitudes across experimental conditions further supports this notion (Figure 6g). We found, however, no significant difference between upregulation and control trials in neither component. Since both pupil-linked arousal (as previously shown^14^) and cortical arousal as indexed by the spectral slope was also not significantly different between upregulation and control trials, these findings likely suggest that these two task conditions may have been associated with similar global arousal states leading to comparable phasic responses to task-relevant target sounds. This heightened arousal state during the control condition may be driven by the rather challenging control task of counting backwards in steps of 7. In summary, we provide first evidence, that through pupil-based self-regulation, we can volitionally modulate the size of the P300 in the context of an attention task. Together with our previous results showing that pupil-based self-regulation systematically modulates the LC-NA system^14^, our findings may additionally support the notion of the LC-NA system being involved in generating the P300^34,45,56^.

We are aware that the interpretation of indirect indices of LC activity warrants caution, especially concerning pupil dynamics. Although the current literature predominantly supports non-luminance-related pupil size fluctuations to be closely linked to activity changes in the LC^10–13^, other mechanisms may be modulating such fluctuations directly or indirectly. Especially arousal-mediating cholinergic brain regions in the basal forebrain^10^, but also dopaminergic nuclei such as the ventral tegmental area and substantia nigra^57^ and hypothalamic orexin neurons^58^ have been shown to be linked to pupil-linked fluctuations. Our previous study combining pupil-BF with (brainstem) fMRI revealed that, even though pupil-BF most consistently modulated activity of the LC, it also led to activity changes in cholinergic and dopaminergic brain regions, albeit less consistently^14^. Similarly, together with the LC-NA system, the dopaminergic system has been proposed to underly the P300 generation observed on the scalp level^52^. Thus, modulatory effects observed in the present study could either be driven via the modulation of NA transmission, leading to the release of other neurotransmitters and/or co-transmitters, or via targeting other neuromodulatory systems directly.

In summary, we provide evidence that acquired pupil self-regulation through pupil-BF training modulates cortical markers of excitability and arousal. Especially the correlation between pupil-linked and cortical measures of arousal supports the interpretation of a shared underlying physiological mechanism. Similarly, the systematic response modulation of the P300 through pupil self-regulation adds to previously reported changes in pupil dilation as a function of pupil-linked arousal states. These findings suggest that pupil-based biofeedback shown to modulate LC activity in humans influences cortical markers of excitability and arousal considered to be fundamental aspects of brain function. Future studies are needed to determine whether pupil-BF could further be used to modulate these aspects in case of disturbances associated with neurological and psychiatric disorders.

## Methods

### General Information

All experimental protocols were approved by the local Cantons Research Ethics Committee of Zurich (KEK-ZH 2018-01078) and performed in accordance with the Declaration of Helsinki. All participants included in the study were healthy adults with normal or contact-lens-corrected vision, and reported no neurological or psychiatric disorders, or medication intake that affects the central nervous system. All participants were asked to avoid caffeine consumption on the day of the testing. During EEG testing sessions (i.e., Experiments 2 and 3), participants sat in a noise-shielded Faraday cage (mrShield^®^ type EEG, CFW EMV-Consulting AG, Reute, Switzerland). All experimental sessions were performed under constant lighting conditions within and across participants. After a detailed explanation of the experimental procedure, all participants gave their informed consent and received financial compensation for their participation (20 CHF per hour). None of the experiments were preregistered.

#### Pupil-based biofeedback

In all experiments reported here, participants were trained using the previously described pupil-BF procedure consisting of three training sessions^14^. Here, we only report the data following these training sessions (Experiments 1 and 3) or from the third training session (Experiment 2). Prior to training, the participants were sufficiently informed about the exact procedure and essential elements of the study. After providing informed consent, participants were presented with possible mental strategies for increasing or decreasing their pupil size and asked to try these mental strategies while using the feedback information displayed on the screen. The suggestions on mental strategies were derived from previous studies in the literature in which the pupil either dilates or constricts depending on different cognitive tasks and mental states^59,60^. Participants’ ocular dominance was then determined (right eye dominance: *n*=10 in Experiment 1; *n*=15 in Experiment 2; *n*=16 in Experiment 3) using a modified Miles technique, as pupil diameter feedback was calculated based on data from the dominant eye only^14^. During the training sessions, participants aimed to acquire appropriate mental strategies to increase (i.e., upregulate) or decrease (i.e., downregulate) their pupil diameter and thus self-regulate their pupil-linked arousal state. In each experimental session, participants performed trials of upregulation (i.e., increase in pupil size) and downregulation (i.e., decrease in pupil size). In Experiment 1 and 3, there was an additional resting (Experiment 1) or counting control (Experiment 3) condition. All trials began with an instruction on the modulation direction of pupil size (either up- or downregulation of pupil diameter) displayed in green (Experiment 2) or pink (Experiment 1 and 3) on a grey background, followed by a baseline measurement. During this baseline phase, participants were presented with a “=” sign above the fixation cross and were instructed to silently count backwards in steps of four (Experiment 1 and 2) or seven (Experiment 3) to maintain their mental state at a constant and controlled level. In the following, participants were applying their mental strategies for pupil diameter up- and downregulation during the modulation phase. The transition from the baseline to the modulation phase was indicated by a change from an equal (”=”) to an “x” sign above the fixation cross. Following modulation, participants received color-coded post-trial feedback on their modulation performance. Therefore, from valid pupil diameter samples, the average across the modulation phase was calculated and the maximum change to baseline was extracted. Average and maximum changes were then displayed on the screen: for successful modulation of pupil size towards the indicated direction (i.e., pupil size during modulation was larger than pupil size during the baseline phase in upregulation trials and smaller than pupil size during the baseline phase in downregulation trials, respectively) the feedback circle indicating the average change was shown in green. For unsuccessful pupil size modulation (i.e., pupil size during modulation was the same or smaller than the baseline pupil size in the upregulation trials or bigger than baseline pupil size in downregulation trials), the circle was displayed in pink (see Figure 2). We accounted for artifacts during pupil self-regulation caused by eye blinks, physiological and measurement-based noise with the following preprocessing steps: (i) rejection of data samples containing physiologically implausible pupil diameter values ranging outside of a pupil size of 1.5 and 9mm – this step ensured that blinks, automatically detected by the eye tracker, were not included in feedback shown to participants; (ii) rejection of physiologically implausible pupil size changes larger than 0.0027mm/s. This previously implemented approach^61,62^ was based on specifications of a study reporting peak velocity of the pupillary light reflex^63^. All colors presented throughout the experiment, were isoluminant to the grey background (RGB [150 150 150]) by determining the relative luminance according to the following combination of red, green and blue components: Y = 0.2126 R + 0.7152 G + 0.0722 B; https://www.w3.org/Graphics/Color/sRGB. Between blocks of trials, participants were given the opportunity to take a self-determined break before moving on to the next block. Throughout the trials of each experiment, participants were asked to fixate their gaze on the fixation dot in the middle of the screen. Stimulus presentation throughout the experiments was controlled using the MATLAB-based presentation software Psychtoolbox 3.0.17. Since the number of trials and trial timings differed slightly for each of the reported experiments, details are reported in each experimental section separately (Figure 2). For further details of the pupil-based biofeedback approach^14^.

#### Statistical analysis

Statistical analyses were performed using IBM SPSS 28 (IBM Corporation, Armonk, NY, USA), R version 4.1.2 (R Core Team), and JASP (version 0.16.2).

All data were tested for normal distribution using Shapiro-Wilk test of normality. For all repeated measures (rm) analyses of variance (ANOVA) models, sphericity was tested using Mauchly’s sphericity test. In case of violation of sphericity, a Greenhouse-Geisser correction was applied. The threshold for statistical significance was set at α=0.05. Correction for multiple comparisons using sequential Bonferroni correction^64^ was applied where appropriate (e.g., post-hoc tests). We used two-tailed tests for all our analysis.

### Experiment 1. Cortical excitability during Pupil-BF

#### Participants

As there are currently no studies showing an influence of pupil-based biofeedback on cortical excitability tested with neurophysiological read-outs, the required sample size was estimated based on a pilot experiment (*n*=7; Cohen’s *d*_z_=1.04), using a power analysis (G*Power version 3.1^65^). It revealed that 15 participants should be included to detect an effect of pupil size self-regulation on MEP amplitude with a two-tailed t-test (difference between upregulation to downregulation condition, normalized to baseline), α=0.05, and 95% power.

We re-recruited nineteen healthy volunteers (14 females, 26±5 years) with normal or corrected-to-normal vision, who previously successfully completed three training sessions of pupil-based biofeedback training^14^ and indicated their willingness to be informed about future studies.

We had to exclude four participants: one because of technical problems with data acquisition, and three because of our analysis exclusion criteria (see *Electromyography data processing and analysis* below). Fifteen participants (11 females, 26±5 years) were included in the final data analysis. One participant was left-handed (Edinburgh Handedness Inventory mean laterality quotient, LQ=-80^66^), one was ambidextrous (LQ=6.67) and thirteen were right-handed (LQ=77.7±18.2). All participants had no identified contraindications for participation according to established TMS exclusion criteria^67–69^ and provided written informed consent. None of the participants reported any major side effects resulting from the stimulation.

#### Pupil-based biofeedback combined with TMS

Participants sat in a comfortable TMS chair with their head resting on a backrest to ensure a stable head position without putting too much strain on the neck of the participants. Participants wore earplugs to protect their hearing and to diminish potential distraction due to the sound of the TMS pulse^69^. We adjusted the position (height) of the screen to the height of the participant^70^. Participants eyes were ∼70cm away from the eye tracker (Tobii Pro Nano, Tobii Technology AG, Stockholm, Sweden) that was positioned below the screen (240P4QPYEB/00, resolution: 1.920×1.200, 61cm; Philips, Amsterdam, Netherlands) to allow for optimal eye tracking and measurement of pupil size during the pupil-based biofeedback procedure as described above. The experiment was coded in MATLAB R2020a (Mathworks Inc., Natick, USA) and presented using the MATLAB-based Psychtoolbox extension 3.0.17^71,72^. Pupil diameter and eye gaze data of both eyes were sampled at 60Hz. At the beginning of the experimental session, the eye tracker was calibrated using a 5-point calibration technique. All participants underwent 2 modulation and 1 control conditions: (i) upregulation, in which they upregulated (i.e., increased) their pupil size, (ii) downregulation, in which they downregulated (i.e., decreased) their pupil size, and (iii) a resting control condition, in which the participants’ task was to stay at rest and let their mind wander freely, without applying modulation strategies. Each condition was repeated twice (i.e., a total of 6 blocks). Each block consisted of 10 trials, which started with a 1s instruction phase indicating the required pupil self-regulation or the control condition, followed by a 4s baseline measurement. Then, the participants applied the trained mental strategies to up- or downregulate pupil size for 15s, or to rest. During this self-regulation or resting phase, two TMS pulses were delivered over their primary motor cortex, where the 1^st^ pulse was applied between 3-4s and 2^nd^ pulse 11s after the modulation onset. Note that for the participant the timing of the 2^nd^ pulse was subjectively variable in relation to the 1^st^ pulse. Cortical excitability was measured by the means of MEPs recorded from the first interosseous muscle (FDI) of the right index finger (see *TMS* and *Electromyography* sections below). After each trial of the modulation condition, participants received post-trial feedback (see Figure 2). Pupil size and ECG were recorded continuously in each trial.

#### TMS Properties

Single-pulse monophasic TMS was delivered using a 70mm figure-of-eight coil connected to the Magstim 200 stimulator (Magstim, UK). The coil was positioned over the hotspot of the FDI muscle. The hotspot was defined as the stimulation site where TMS delivery resulted in the most consistent and largest MEPs in the resting muscle. The coil was held tangential to the surface of the scalp, with the handle pointing backward and laterally at 45° away from the nasion-inion mid-sagittal line, resulting in a posterior-anterior direction of current flow in the brain. Such a coil orientation is thought to be optimal for inducing the electric field perpendicular to the central sulcus resulting in the stimulation of primary motor cortex neurons^73,74^. The optimal coil location was marked with a semi-permanent marker on the head and the coil orientation was additionally indicated by a line drawn along the edge of the coil on a piece of tape on the participant’s head. This ensured that the center of the coil was kept over the determined hotspot, and that the coil orientation was consistent throughout the experiment. At the beginning of the experiment, for each participant, we determined the resting motor threshold (RMT), defined as the lowest TMS intensity to elicit MEPs with peak-to-peak amplitude ≥0.05mV in the relaxed muscle, in five out of 10 consecutive trials, i.e., 50% (^75^ mean RMT=41.7 ± 7.3% MSO, range: 29–54%). We then applied single-pulse TMS with an intensity of 120% of the individual RMT during the experiment (50.8±8.8% MSO, range: 35–64%). Two TMS pulses were delivered during the pupil modulation phase in the upregulation, downregulation or resting control conditions (see Figure 3). The first pulse was jittered between 3-4s and the second pulse and the second pulse 11s after modulation (or rest) onset to avoid TMS pulse anticipation. Two pulses delivered per trial resulted in a sample of 40 MEPs collected per condition.

#### Electromyography

The muscle response was recorded by a surface electromyography (EMG) electrode (Bagnoli™ DE-2.1 EMG Sensors, Delsys, Inc.) placed over the right FDI muscle. Raw signals were amplified (sampling rate, 5kHz), digitized with a CED micro 1401 AD converter and Signal software V2.13 (both Cambridge Electronic Design, Cambridge, UK), and stored on a personal computer for off-line analysis. The timing of the TMS pulses and EMG data recording was controlled by MATLAB R2020a (MathWorks, Inc., Natick, MA, USA) connected to the CED via a custom microcontroller. Muscular relaxation was constantly monitored through visual feedback of EMG activity and participants were instructed to relax their muscles if necessary.

#### Electrocardiography

The cardiac data were recorded with a wireless, Bluetooth-based, wearable Shimmer3 system (© Shimmer 2017^TM^, Realtime Technologies Ltd)^76,77^ and the accompanying Shimmer MATLAB^TM^ Instrument Driver 2.8a available at https://github.com/ShimmerEngineering/Shimmer-MATLAB-ID. ECG was recorded separately in each trial and sampled at 512Hz from five electrodes (EL503 Ag/AgCl Pre-Gelled Vinyl 1-3/8’’ Electrodes, Biopac Systems, Inc.), attached under the right collarbone, under the left collarbone, to the right lower rib, to the left lower rib and to the mid-left upper rib, as suggested by the Shimmer Manual. For subsequent offline analyses, the ECG biphasic signal recorded by the electrodes positioned over the left lower rib and under the right collarbone was used.

#### Pupil data offline processing and analysis

(Pre-)processing of the pupil data was conducted using MATLAB R2020a (MathWorks, Inc., Natick, MA, USA). Recorded pupil size (in mm) and gaze data were visually inspected to ensure that participants followed the instructions to look at the fixation dot in the center of the screen during the baseline and modulation phase of the experiment. Then, pupil data of both eyes were systematically preprocessed using the guidelines and standardized open source pipeline published by Kret and Sjak-Shie^78^. Invalid pupil diameter samples representing dilation speed outliers and large deviations from trend line pupil size (repeated four times in a multipass approach) were removed using a median absolute deviation (MAD; multiplier in preprocessing set to 12 – a robust and outlier resilient data dispersion metric^78^). Further, temporally isolated samples with a maximum width of 50ms that border a gap larger than 40ms were removed. Then, mean pupil size time series were generated from the two eyes which were used for all further analyses. The data were resampled with interpolation to 1000Hz and smoothed using a zero-phase low-pass filter with a recommended cutoff frequency of four^78^. Finally, preprocessed pupil diameter was corrected relative to baseline, by computing mean pupil size of the last 1000ms before the start of the modulation phase of each trial and subtracting this value from each sample of the respective modulation phase, as previously recommended^79^.We quantified the difference between pupil diameter in the up-, downregulation and resting control conditions by calculating the average of baseline-corrected pupil diameter measured for 150ms before each TMS pulse, to exclude the influence of the TMS-pulse itself on the pupil diameter. We then compared the averaged pupil diameters with a rmANOVA with the factor condition (upregulation, downregulation, and rest).

#### Electromyography data processing and analysis

(Pre-)processing of the EMG data was conducted using MATLAB R2020a (MathWorks, Inc., Natick, MA, USA). The EMG data was band-pass filtered (30-800Hz, notch filter = 50Hz). Filtering was applied separately for the pre-TMS background EMG (bgEMG; measured for 100ms between 105ms and 5ms before the TMS pulse) and post-TMS period containing peak-to-peak MEP amplitude to avoid “smearing” the MEP into bgEMG data. The peak-to-peak amplitude was defined as the peak-to-peak amplitude between 15 to 60ms after the TMS pulse. Trials with root mean square bgEMG above 0.01mV were removed from further analyses to control for unwanted bgEMG activity^80,81^. For the remaining trials, the mean and standard deviation of the bgEMG was calculated for each participant. Trials with bgEMG > mean + 2.5 standard deviations were also excluded. Data from participants who showed differences in root mean square bgEMG>0.001mV between pupil size upregulation and downregulation conditions were removed to avoid the possibility that the differences in bgEMG could drive the effects on MEP amplitude (note, that this exclusion criterion is especially conservative, as bgEMG itself could be influenced by successfully modulated arousal level). Based on this criterium, we removed data from 3 participants.

MEP amplitudes from the modulation conditions (up- and downregulation) were normalized to the resting control condition and tested using rmANOVA with the factor condition (upregulation and downregulation, both normalized to rest). We additionally analyzed the effects with bgEMG added as covariate. BgEMG was calculated as (bgEMG upregulation-bgEMG downregulation)/average(bgEMG upregulation, bgEMG downregulation). We controlled for the bgEMG to make sure that the effects of the pupil self-regulation were not driven by elevated bgEMG in one of the conditions. Additionally, we performed a Bayesian rmANOVA with the factor condition (upregulation, downregulation, rest) on bgEMG to evaluate the absence versus presence of a significant difference in bgEMG between self-regulation and control conditions.

Further, since participants applied mental strategies from different categories (i.e., emotional versus activation-related) to upregulate the pupil size, we split the data set into two groups using (i) purely emotional strategies and (ii) a mix of activation and emotional strategies, respectively. We wanted to disentangle whether the effects on cortical excitability are not driven by motor imagery shown to activate the primary motor cortex^42,43^. The differences between the upregulation and downregulation conditions in each group were tested with two-tailed paired samples t-tests and corrected for multiple comparisons using sequential Bonferroni correction^64^.

#### Cardiac data (pre-)processing and analyses

ECG R peaks were automatically detected and, if necessary, manually corrected after visual inspection using the MATLAB-based toolbox PhysioZoo^82^. Data segments consisting of non-detectable peaks or poor quality were excluded from further analyses. Resulting R-R intervals for which both R peaks are occurring in the baseline and modulation (or rest) phase were extracted and further processed in MATLAB (R2020a).

Heart rate, reflecting cardiovascular dynamics controlled by an interplay between the sympathetic and parasympathetic nervous system, was calculated by dividing 60 through the respective R-R intervals of the modulation or rest (control) phase. Furthermore, we computed the root mean square of successive differences (RMSSD) calculated based on R-R-intervals. We chose RMSSD because it is relatively free from breathing influences^83^ and can be computed for intervals as short as 10s^84,85^. It represents a beat-to-beat measure of HRV in the time domain that is assumed to give meaningful insights about parasympathetic activity of the autonomic nervous system. Heart rate and RMSSD calculated for each modulation phase were averaged across the respective conditions. Heart rate and HRV (i.e., RMSSD) differences between pupil size up- and downregulation (normalized to the resting control condition) were tested using two-tailed paired samples t-tests.

#### Link between pupil-based, cortical, and cardiovascular arousal

To investigate the relationship between pupil-based self-regulation and differences in cortical (i.e., MEP) arousal markers, we subjected the different modulation indices and MEP normalized to the rest control condition to a Pearson correlation analysis. To investigate the relationship between differences in cortical (i.e., MEP) and cardiovascular arousal markers (heart rate, HRV), we subjected the MEP, heart rate and HRV (normalized to the rest control condition) to a Pearson correlation analysis.

### Experiment 2. Cortical arousal during Pupil-BF

#### Participants

A total of 25 eligible participants (13 females; 27±6 years) were recruited for this study from an in-laboratory pool of former participants^14^ or via an online advertisement. The sample size was similar to^28^ or larger than^27^ in previous experiments investigating the spectral slope as an electrophysiological marker of arousal levels in humans. Exclusion for personal reasons (*n*=1) and difficulties in adhering to the study procedures in the first two sessions (*n*=1) resulted in a final sample size of 23 participants (12 females; 27±5 years) reported in Experiment 2.

#### Pupillometry recordings during pupil-based biofeedback

During all sessions, participants sat in a Faraday cage (mrShield^®^ type EEG, CFW EMV-Consulting AG, Reute, Switzerland) with their chins on a chin rest adjusted to a standardized height. The height of the chair was individually adjusted to suit the participants. The chin rest was positioned at a standardized distance of 0.65m between the participants’ eyes and the eye tracker to ensure optimal eye tracking and pupil diameter measurement. The experiment was coded in MATLAB R2013b (Mathworks Inc., Natick, USA) and presented using the MATLAB-based Psychtoolbox extension 3.0.17^71,72^. Throughout the experiment, pupil size and gaze of both eyes were measured using an infrared eye tracker (Tobii tx300, Tobii Technology AG, Stockholm, Sweden) with a sampling rate of 60Hz. At the beginning of each experimental session, the eye tracker was calibrated using a 5-point calibration technique.

Here, we only report the data of training day 3 of pupil-based biofeedback (not reported before) where we simultaneously recorded EEG and ECG data. In each of the training sessions, participants performed 60 trials in total, separated in three upregulation and three downregulation blocks, each consisting of 10 trials. All trials began with a 2s instruction phase on the modulation direction of pupil size (either up- or downregulation of pupil size), followed by a 3s baseline measurement. In the following, participants were applying their mental strategies for pupil diameter up- and downregulation during the modulation phase. As compared to the training procedure described in^14^ a longer modulation phase of 30s was introduced in this session due to additional blood pressure measurements not reported here. However, similar to our previous study, we will report results of the first 15s of modulation. Following modulation, participants received post-trial feedback on their modulation performance for 2s. A short pause of 2s separated each trial.

#### Electroencephalography Recordings

The EEG signal during pupil-BF was recorded with 64 gel-based Ag/AgCl active surface electrodes (Brain Products, Munich, Germany), an actiCHamp Plus amplifier and the multifunctional, proprietary software BrainVision Recorder software (Brain Products, Munich, Germany) at a frequency of 1kHz. Using the actiCAP SNAP holders (Brain Products, Munich, Germany), the electrodes were placed in the EEG cap according to the standardized International 10-20 System. After assembly on the participants, all the electrodes were filled with a conductive saline electrode gel using a syringe. Circular movements during the filling process helped to lower the impedances between the electrodes and the scalp. Impedances were kept below 20 kOhm.

#### Electrocardiography

To acquire cardiac data, the BIOPAC MP160 system was used together with the provided AcqKnowledge^®^ software (BIOPAC^®^ Systems, Inc., Goleta, USA). Disposable surface electrodes (all-purpose electrodes, BIOPAC^®^ Systems Inc., Goleta, USA) were used. Continuous ECG data were collected at 1kHz from two surface electrodes, one under the right clavicle and one on the lower left rib. A reference electrode was placed under the left collarbone. Before mounting the ECG electrodes, participants’ skin was prepared with fine sandpaper and disinfected for optimal impedance. An impedance test and visual inspection ensured good signal quality before the start of the experiment. Additionally, continuous blood pressure and respiration effort was measured throughout the experiment, however the data are not reported here.

#### Offline (Pre-)Processing and Statistical Analyses of Pupil Data

All data (pre-)processing steps and statistical analyses on the pupillary data were performed using MATLAB R2018a, R2022a (Mathworks Inc., Natick, USA), SPM 1D Toolbox (SPM1D M.0.4.8; https://spm1d.org/), and SPSS version 28.0.1.1. (IBM Corp., Armonk, NY, USA). Visual inspection of recorded pupil size (in mm), gaze data, and video recordings ensured that participants followed instructions to look at the fixation cross in the center of the screen during the baseline and modulation phases and did not use eye movement-related strategies to modulate their pupil diameter (e.g., squinting or large vergence movements). Trials containing systematic squinting or vergence movements of the eyes (deviations in visual angle greater than ∼16° on the x- and ∼10° on the y-axis) that were at risk of biasing the effective pupil size^70^ were excluded from further analysis.

Systematical preprocessing steps of the pupillary data followed the standardized open-source pipeline^78^ as described in Experiment 1. Following preprocessing and the removal of trials with more than 50% missing data point (excluded: upregulation = 3%; downregulation = 6%; *z*=1.33; *p=*0.14; *r*=0.28) across the baseline and modulation phases, baseline-corrected mean pupil size time series of the modulation phase were averaged across the up- and downregulation conditions for each participant. Further, baseline pupil diameter used for trial-based baseline correction was calculated for each up- and downregulation condition. To draw conclusions on whether participants volitionally self-regulate their pupil size on day 3 of pupil-BF training, baseline-corrected pupil values were used to compute a pupil modulation index. This modulation index assesses the average difference between baseline-corrected pupil sizes during upregulation and downregulation (i.e., up-down) over all *n*=15’000 data points in the 15s modulation interval: Since successful upregulation is reflected in positive baseline-corrected pupil size and successful downregulation in negative baseline-corrected values, larger modulation indices correspond to better condition-specific modulation performance.

To ensure stable pupil size during baseline periods, the absolute pupil data of both conditions of the last second of the baseline were assessed using a paired samples t-test. To examine whether participants were able to self-regulate pupil size during the reported session, baseline-corrected pupil size averaged across the upregulation and downregulation conditions, respectively were compare using a paired samples t-test. Further, we averaged the modulation index time series (i.e., up-down) across the 15s modulation phase and conducted a one-sample t-test to investigate whether the indices differed significantly from zero.

#### Offline (Pre-)Processing and Statistical Analyses of EEG Data

All EEG data analyses were performed using the EEGLAB Toolbox^86^ and according to a standardized Automagic pipeline^87^ running on MATLAB R2022a (Mathworks Inc., Natick,USA) and the spectral parameterization toolbox FOOOF^88^ in combination with custom codes written in MATLAB. To account for human variability during (pre-)processing, the (pre-)processing steps of the EEG dataset were semi-automated by adhering to the open-source Automagic pipeline^87^ and standardization to pre-set thresholds was strictly applied throughout the process. First, the raw data were down-sampled to a frequency of 250Hz and high- and low-pass filters were applied (i.e., high pass filter = 0.1Hz; low pass filter = 45Hz) to minimize slow drifts and trends and to eliminate 50Hz line noise.

The EEGLAB plugin clean_rawdata enabled automatic detection and separation of flatline and noisy channels and time windows from EEG activity^89^. The artifactual data parts and channels were plotted for visual inspection. Discarded channels were interpolated and the resulting scalp EEG data re-referenced to the average electrical activity of all channels present, according to the Common Average Reference^90^. Then, Independent Component Analysis (ICA) was applied using the open-source Multiple Artifact Rejection Algorithm (MARA) implemented in the EEGLAB toolbox. The machine learning algorithm automatically marked and separated multiple types of artifacts, such as eye movements, muscle, and cardiac activity from brain components with a corresponding estimated probability of belonging to that respective category. All components were plotted for visual inspection along their assigned component category and only components labelled as brain or other with a likelihood of >85% were included in all further analyses. Following these preprocessing steps, the EEG data were divided into up- and downregulation epochs from -3 (i.e., baseline phase) to 15s after modulation onset (i.e., modulation). All datasets of each trial were then divided into 3s time bins, resulting in a baseline bin and 5 bins for the modulation phase. Time bins containing artifacts as previously marked by the EEGLAB plugin clean_rawdata, were excluded from further analyses (Bad Epochs Upregulation = 12%, Bad Epochs Downregulation = 11%; paired samples t-test: *z*=-0.10; *p*=.92; r=-0.02). Next, we calculated the power spectral densities (PSD; i.e., the power distribution in the frequency domain) using Welch’s method implemented in the EEGLAB toolbox for different time bins, channels, and self-regulation conditions with a frequency resolution of 1Hz for 1 to 43Hz with an overlap of 50% and averaged across trials leading to an average power spectrum for both the up- and downregulation conditions per time window for each participant. To estimate the aperiodic component of the power spectra (i.e., spectral slope) we used the spectral parameterization toolbox FOOOF with the FOOOF MATLAB wrapper and the default settings, except setting the maximum number of peaks to 3. Based on previous studies, the frequency range was set to 30 to 43Hz (to ensure that no frequencies affected by the low-pass filter were included) which has been shown to reliably differentiate different arousal states in humans^27^ and E/I changes in animal work^26,27,29^. In a secondary analysis, we estimated the parameters for a larger frequency range of 2 to 40Hz^29^, which is reported in Supplementary Figure 4. Because recent literature suggests that arousal state-induced fluctuations in spectral slope are not localized but rather are reflected broadly across the scalp, spectral slopes for each participant were analyzed as a whole-brain average, as opposed to conducting analyses on individual channels^27,29^. After a first fitting the original power spectrum as a full model (i.e., as a combination of aperiodic and periodic, oscillatory components), a linear regression line enabled the extraction of the aperiodic parameters (i.e., the slope, the error and the fit of the model, thus R^2^).

Statistical analyses on the EEG data were performed using SPSS, version 28.0.1.1. (IBM Corp., Armonk, NY, USA). Even though the FOOOF algorithm estimates both periodic (i.e., oscillatory) and aperiodic components (i.e., the slope) of the power spectrum, we only performed statistical analysis of the spectral slope as an indicator of the cortical arousal state of the brain. First, to examine the potential effects of pupil self-regulation on the spectral slope, we extracted the estimated slope for each time bin of the modulation phase and averaged these estimates across the times bins of the up- and downregulation condition, respectively. Then, we subjected the spectral slopes to a two-sided paired samples t-test. Next, to investigate whether effects of pupil self-regulation on spectral slope estimates evolved over time, we subjected the slope estimates to a rmANOVA with the factors time bin (baseline versus the five different time bins of the modulation) and condition (upregulation versus downregulation). In case of significant effects, post-hoc analyses were conducted and corrected for multiple comparisons using sequential Bonferroni corrections^64^. Analogously to the pupil modulation index, we calculated a similar score for the spectral slope (i.e., slope_up_ – slope_down_), reflecting the differences in the steepness as a function of the modulation conditions. Hence, the higher the modulation index, the greater the differences in the spectral slope between the two modulation conditions.

#### Offline (Pre-)Processing and Statistical Analyses of Cardiac Data

ECG data were (pre-)processed using MATLAB R2022b (MathWorks, Inc., Nattick, MA, USA) and the open-source MATLAB based software PhysioZoo^82^ as described in Experiment 1. Technical problems during data saving led to the exclusion of one participant resulting in *n*=22. Based on the valid R-R-intervals, heart rate (in BpM) and HRV values (i.e., RMSSD) were extracted and averaged across the respective condition (i.e., upregulation and downregulation). To compare heart rate between up- and downregulation conditions, we conducted a two-sided paired samples t-test. Since RMSSD values showed a significant deviation from normal distribution (Shapiro-Wilk test *p* <.05), condition differences were examined using a Wilcoxon Signed Rank test. Analogously to the pupil modulation index, we calculated similar scores for heart rate and RMSSD data (i.e., heart rate_up_ - heart rate_down_; RMSSD_down_ – RMSSD_up_). The higher the modulation indices, the greater the differences in heart rate and RMSSD between the two modulation conditions.

#### Link between pupil-based, cortical, and cardiovascular arousal

To investigate the relationship between pupil-based self-regulation and differences in cortical (i.e., spectral slope) arousal markers, we subjected the different modulation indices to a Pearson correlation analysis. Similarly, we inspected the relationship between cortical (spectral) and cardiovascular arousal markers (i.e., heart rate and RMSSD) via the computation of Pearson correlation coefficients (i.e., heart rate) and Spearman rho coefficients (i.e., RMSSD; due to a deviation from normal distribution (i.e., Shapiro-Wilk test, *p* < .05)).

### Experiment 3. Cortical arousal and the P300 during pupil-BF and a simultaneous auditory oddball task

Pupil size and reaction time data during pupil-BF combined with an auditory oddball task has been analyzed and reported before^14^. Here, we focus on reporting the EEG data acquired in the same study.

#### Participants

22 participants (15 females; 26 ± 6 years) participants trained with pupil-BF^14^ performed the oddball task while simultaneously self-regulating pupil size. Sample size was similar to or larger than previous investigations of evoked EEG and pupillary responses^40,41^. In addition to the two participants that had to be excluded for pupil size measurements, an additional EEG dataset needed to be excluded due to a recording issue leading to less than 50% of the trials leaving a sample size of *n*=19 participants (12 females; 26±6 years). If the last session of pupil-BF occurred more than 10 days before the scheduled pupil-BF oddball experiment, participants received one session of re-training (i.e., the same trial routine as described for day 3).

#### Pupil-based biofeedback combined with an auditory oddball task

The auditory oddball paradigm during pupil-BF consisted of 186 target (∼20%) and 750 standard (∼80%) tones (pitch of 440 and 660Hz counterbalanced across participants) presented for 100ms via headphones at ∼60dB (Panasonic RP-GJE125, Panasonic). The oddball task was embedded in the pupil size up- and downregulation phases of the pupil-BF paradigm (Figure 2). In addition to the self-regulation conditions, a third control condition was introduced in which participants were instructed to simply count backwards in increments of seven without any pupil modulation occurring. After every 6^th^ counting trial, participants were asked to enter the final number of the control condition to ensure that they completed this task. The rationale behind the counting condition was to control for effort effects related to pupillary self-regulation and to examine differences in oddball responses between self-regulation and natural fluctuations in pupil size. Before each task, participants were reminded to focus on their acquired mental strategies while equally paying attention to the tones being played. After an instruction on the self-regulation direction (upregulation or downregulation) or the counting control condition at the start of each pupil-BF oddball trial, participants underwent a baseline measurement of 4s and were asked to silently count backwards in increments of seven. In the following, participants applied their acquired mental strategies to either up- or downregulate their pupil size for a period of 18s or continued counting. During this self-regulation or counting period, eight tones (i.e., one or two target tones and seven or six standard tones, respectively) were presented while they were asked to respond to the targets as quickly as possible by pressing a key. The first tone (jittered 1.8-2.2s after the baseline phase) was always a standard tone, ensuring that participants were given sufficient time to modulate their pupil size before the first target appeared. The inter-stimulus interval (ISI) between tones varied randomly between 1.8–2.2s. At least two standard tones were presented between two targets, resulting in a minimum ISI of 5.4s between two targets. After each self-regulation trial, participants received 2s of post-trial feedback on their average and maximum change in pupil diameter and were given a short 2s break, before the next trial began. In the counting control trials without feedback phases, counting was followed by a 4s break to ensure comparability of the trial times. The pupil-BF self-regulation conditions were presented in nine blocks (i.e., three per condition), with each block containing 13 upregulation, downregulation, or control trials. The blocks occurred in pseudorandomized order as previously described. The number and order of tones were kept congruent for each pupil-BF condition.

#### Physiological Data Acquisition

Eye tracking, pupillometry, and EEG data were acquired throughout the experiment in a Faraday cage as described in Experiment 2. ECG and respiratory data were acquired simultaneously but are not reported here.

#### Offline (Pre-)Processing and Statistical Analyses of Pupil Data

Baseline-corrected pupil upregulation, downregulation, and control time series and pupil dilation responses towards target and standards have been previously reported^14^ and are not reported here.

#### Offline (Pre-)Processing and Statistical Analyses of EEG Data

Preprocessing of EEG data was performed as described in Experiment 2. Following preprocessing, the EEG data were divided into epochs around standard and target tones from -500ms to 2000ms after tone onset for upregulation, downregulation, and control trials.

##### Spectral slope analysis

For the estimation of the spectral slope changes evoked by the tone presentation, all datasets of each trial were divided into 500ms time bins, resulting in a baseline bin (i.e., -500-0ms) and four time bins following tone presentation (0-500; 500-1000; 1000-1500; 1500-2000ms). Time bins containing artifacts as previously marked by the EEGLAB plugin clean_rawdata, were excluded from further analyses (Targets Bad Epochs: Up = 8%, Down = 7%, Control = 10%; Friedman ANOVA: χ^2^=0.57; *p=*.75; W=0.06 ; Standards Bad Epochs: Up = 9%, Bad Epochs = 10%, Bad Epochs = 10%; Friedman ANOVA: χ^2^=1.13; *p=*.57; W=0.11). Similar to Experiment 2, we used the Welch’s method to calculate the PSD and used the FOOOF toolbox to estimate the spectral slope. Here, we set the frequency range to 30 to 43Hz, which has been shown to most consistently track attention-related task performance^28^. Here, spectral slopes for each participant were analyzed both as a whole-brain average (i.e., averaged across channels) as well as on individual channel level (i.e., via cluster-based permutation tests) to investigate whether there are topographic differences in spectral slope during prior to and following tone onset. After first fitting the original power spectrum as a full model (i.e., as a combination of aperiodic and periodic, oscillatory components), a linear regression line enabled the extraction of the aperiodic parameters (i.e., the slope, error and the fit of the model, thus R^2^). Then, we extracted the estimated slope for each time bin, channel, tone, and self-regulation condition. Statistical analyses on the spectral slope averaged across channels were performed using SPSS, version 28.0.1.1. (IBM Corp., Armonk, NY, USA). To investigate whether effects of pupil self-regulation and tone presentation on spectral slope estimates evolved over time, we subjected the slope estimates to a repeated measures ANOVA with the factors time bin (Baseline versus bin 2 versus bin 3 versus bin 4 versus bin 5), self-regulation condition (upregulation versus downregulation versus control) and tone condition (standard versus target). In case of significant effects, post-hoc analyses were conducted and corrected for multiple comparisons using sequential Bonferroni corrections^64^.

##### ERP analysis

For the calculation of the ERPs, the previously defined epochs were further cut to -100 to 800ms relative to tone onset. Epochs containing artifacts as previously marked by the EEGLAB plugin clean_rawdata or epochs with an incorrect behavioral response (false alarm, missed trial) were excluded from further analyses (Targets: Bad Epochs_Up_ = 14%, Bad Epochs_Down_ = 12%, Bad Epochs_Control_ = 16%; Standards: Bad Epochs_Up_ = 10%, Bad Epochs_Down_ = 11%, Bad Epochs_Control_ = 12%; rmANOVA, sound*condition interaction: *F*_(2,36)_=3.13, *p*=.06). This led to exclusion of >50% of trials in most conditions for two participants. We report analyses without these participants (*n*=17) in Supplementary Figure 5. Epochs were baseline corrected by subtracting the average amplitude during the period prior to tone onset (- 100 – 0ms). Visual observation of the scalp topographies suggested a widely distributed difference between conditions, so we used cluster-based permutation t-tests to analyse the P300 across the full scalp. This method controls for multiple comparisons across space and time. Non-parametric cluster-based permutation t-tests^91^ were implemented in the Fieldtrip toolbox (version 2022.12.12)^92^ in MATLAB R2022a. Condition labels were randomly shuffled to obtain the permutation distribution (5000 permutations, Monte-Carlo method), which was compared to the actual distribution to estimate significance (α=0.05, two-sided). P-values were corrected for multiple comparisons using sequential Bonferroni correction^64^.Spatiotemporal clusters were calculated with paired t-tests for downregulation versus upregulation, downregulation versus cognitive control, and upregulation versus cognitive control. For the assessment of the P300, we defined a time window from 252 – 600ms after tone onset; for the N100 we used 52 – 200ms after tone onset. To assess the specificity of the pupil self-regulation modulation effect on the target-evoked P300, we assessed differences between P300 in response to standard and target sounds with a 2*3 rmANOVA (factor sound (targe versus standard), factor condition (upregulation versus downregulation versus control). To this end, we extracted the mean amplitude in the 252–600ms time window from channels showing a prominent difference between up- and downregulation (C3, CP5, CP1, Pz, P3, P4, P8, CP6, CP2, C4, C1, C5, CP3, P1, P5, PO7, PO3, PO4, P6, P2, CPz, CP4, C2, Cz).

##### Correlation between P300 and spectral slope

To assess the correlation between the spectral slope and P300, condition averages for both measures were Z-scored. We used the R package rmcorr (0.6.0)^93^ to compute the repeated measures correlation coefficient across modulation conditions.

## Data availability

Processed pupil, MEP, EEG, heart rate, and HRV data will be made openly available on the ETH Library Research Collection.

## Acknowledgments

The authors would like to express their gratefulness to all participants of the study. We further thank Stephanie Huwiler and the members of the Hochschulmedizin Zürich Flagship project STRESS for fruitful discussions. S.N.M. is supported by the Hochschulmedizin Zürich Flagship project STRESS and the ETH Career Seed Award. N.W. is supported by the National Research Foundation, Prime Minister’s Office, Singapore under its Campus for Research Excellence and Technological Enterprise (CREATE) programme and funded by the SNSF and Innosuisse BRIDGE Discovery grant (40B2-0_203606). The funders had no role in study design, data collection and analysis, decision to publish or preparation of the manuscript.

## Author Contributions

S.M., W.P.S., M.B., N.W., and S.N.M. were involved in conceptualization and design of the study, S.M., W.P.S., B.B., and S.N.M. acquired the data, M.W., S.M., W.P.S., M.B., M.C., N.W., and S.N.M. planned the analysis, M.W., W.P.S., S.M., and S.N.M., analyzed the data; M.W., W.P.S., S.M., N.W., and S.N.M. interpreted the data, M.W., S.M., W.P.S., and S.N.M drafted the manuscript and M.B., B.B., M.C. and N.W. substantively revised it.

