## Supplementary material for "Pupil self-regulation modulates markers of cortical excitability and cortical arousal"

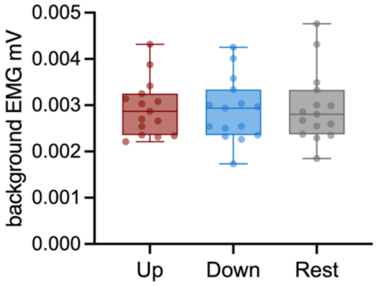


**Supplementary Figure 1. Background EMG.** The root mean square of the background EMG activity (measured for 100ms between 105ms and 5ms before the TMS pulse) did indeed not differ between conditions (Bayesian rmANOVA with the factor condition [upregulation, downregulation and control]: $BF_{01}$=4.83, i.e., moderate evidence for the H0).

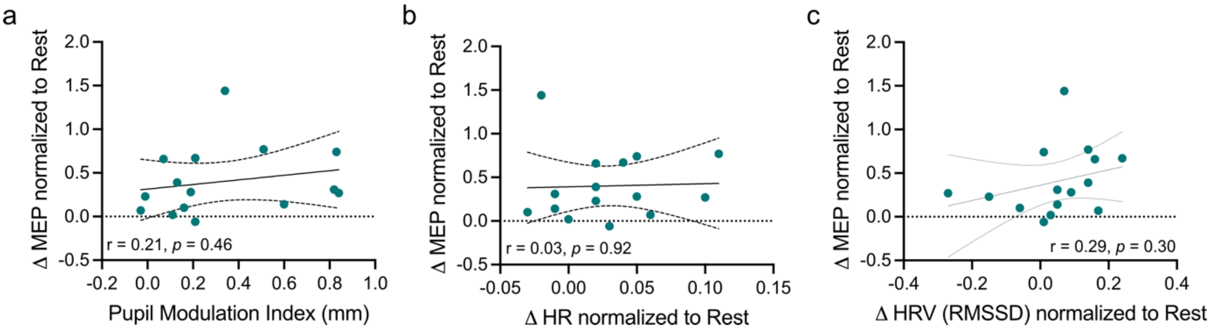


**Supplementary Figure 2. Link of pupil, heart rate, and heart rate variability modulation with MEP.** (a) Non-significant Pearson correlation coefficient between pupil modulation indices (i.e., the difference between pupil size changes in up- versus downregulation trials, Up-Down, calculating the average of baseline-corrected pupil diameter measured for 150ms before each TMS pulse, to exclude the influence of the TMS-pulse itself on the pupil diameter) and differences in MEP between up- and downregulation trials, Up-Down, normalized to control rest condition. (b) Non-significant Pearson correlation coefficient between differences in MEPs (upregulation-downregulation, normalized to control rest condition) and differences in heart rate (upregulation-downregulation, normalized to control rest condition). (b) Non-significant Pearson correlation coefficient between differences in MEPs (upregulation-downregulation, normalized to control rest condition) and differences in heart rate variability (RMSSD, downregulation- upregulation, normalized to control rest condition). Dots indicate individual participants.

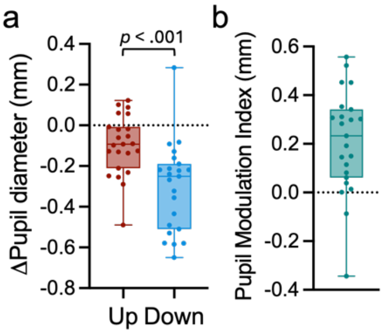


**Supplementary Figure 3. Pupil self-regulation data for 30s time windows.** (a) Baseline-corrected pupil up- and downregulation averaged across the 30s modulation phase for each condition as compared to baseline pupil size in the last second before modulation. Similar to the 15s modulation window, the significant difference between up- and downregulation (two-sided paired samples t-test: $t_{(22)}$=4.73, $p$<.001) were mainly driven by strong downregulation ($t_{(22)}$=-6.75, $p$<.001). (b) Baseline-corrected pupil modulation index reflecting the difference between upregulation and downregulation time windows averaged across the 30s modulation phase. Boxplots indicate median (center), 25th and 75th percentiles (box), maximum and minimum values (whiskers).

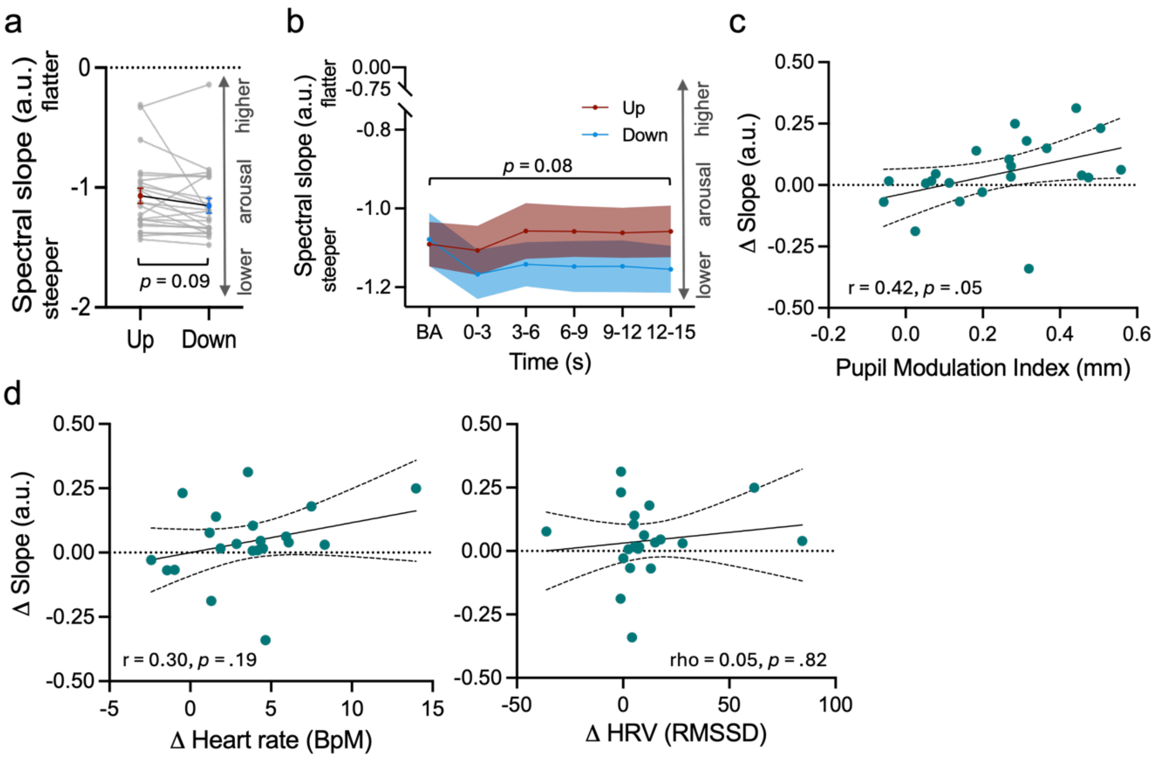


**Supplementary Figure 4**. **Spectral slope during pupil self-regulation extracted for 2-40Hz.** (a) The spectral slope estimated for 2-40 Hz averaged across all channels and time windows of pupil size up- (red) and downregulation (blue), indicating steeper slopes during down- as compared to upregulation (trend-level: $t_{(22)}$=1.8; $p$=.09). (b) The spectral slope estimated for 2-40 Hz averaged across all channels of up- (red) and downregulation (blue) trials for time bins of 3s during baseline (i.e., -3-0s) and the modulation phase (0-3s, 3-6s, 6-9s, 9-12s, 12-15s). Similar to the 30-43Hz, we found a trend for an interaction between time bins and self-regulation condition ($F_{(1.78, 39,14)}$=2.84, $p$=.08). (c) Pearson correlation coefficient between pupil modulation indices (i.e., the difference between pupil size changes in up- versus downregulation trials, Up-Down) and differences in spectral slope between up- and downregulation trials averaged across the 15s modulation phase, indicating a linear relationship (i.e., trend-level) between pupillary and cortical markers of arousal. (d) Non-significant Pearson (left panel) and Spearman rho correlation coefficients (right panel) between differences in spectral slope (upregulation-downregulation) and differences in heart rate (left panel; upregulation-downregulation) and RMSSD (right panel; downregulation-upregulation). Shaded areas indicate s.e.m. Dots indicate individual participants.

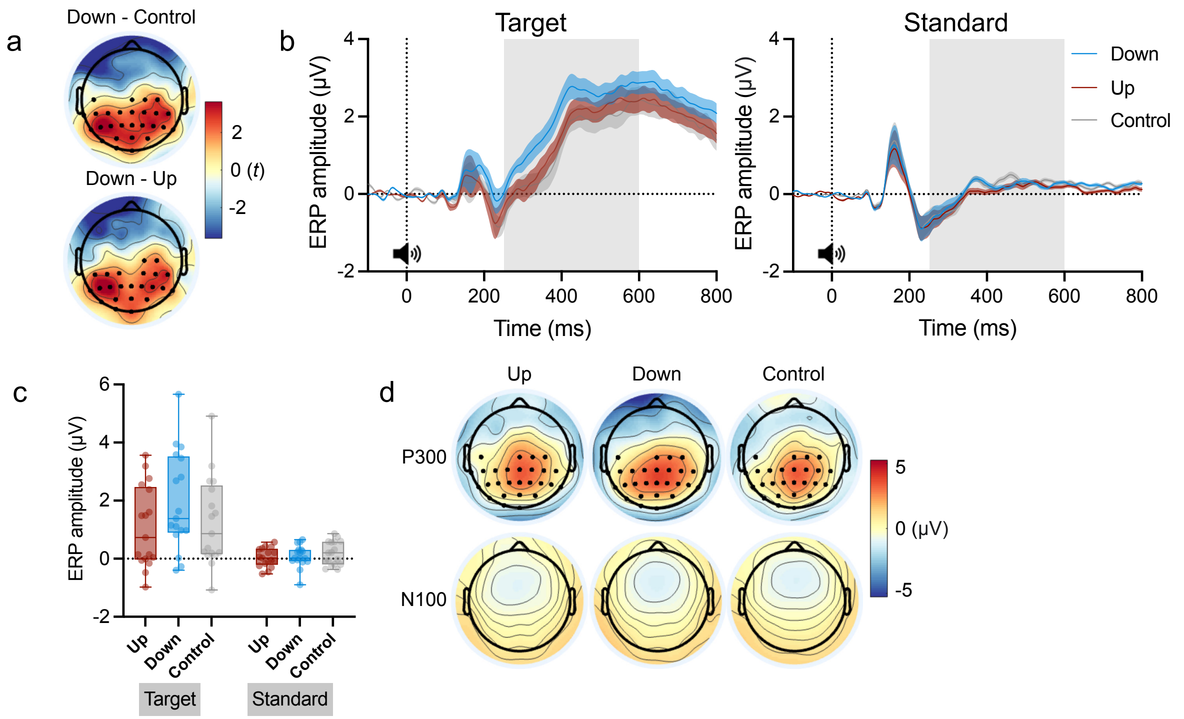


**Supplementary Figure 5: Effects of pupil self-regulation on the P300 and N100 during the oddball task for 17 participants.** Two participants were excluded because >50% of epochs had to be excluded after cleaning the raw data. (a) Resulting t-values of the cluster-based permutation t-tests. Similar to the results reported in the main text (Figure 6c-e), the P300 differed significantly between the downregulation and cognitive control condition (cluster-based permutation t-test on ERPs; 252–600ms after target sound onset: $t$=1796.45, $p$=.0024). This difference was most pronounced from 252–440ms after sound onset in a broad centroparietal cluster (CP5, CP1, Pz, P3, P7, O1, Oz, P4, P8, CP6, CP2, C4, C5, TP7, CP3, P1, P5, PO7, PO3, POz, PO4, P6, P2, CPz, CP4, C2, Cz). There was also a significant difference between downregulation and upregulation in a broad centroparietal cluster from 252 – 388 ms after stimulus onset (t=1184.97, p=.0094; CP5, CP1, Pz, P3, P7, O1, Oz, P4, P8, CP6, CP2, C4, CP3, P1, P5, PO7, PO3, POz, PO4, P6, P2, CP4, C2). We did not observe a significant difference between the upregulation and cognitive control conditions. Most prominent channels are marked with a black dot. (b) ERP time series averaged across the most prominent channels, resulting from the cluster-based permutation t-test between Downregulation and Control. The grey box indicates the time window (252-600ms) for which the P300 was compared between the modulation conditions. (c) A repeated measures ANOVA on the P300 shows a significant main effect of sound ($F_{(1,16)}$=20.71, $p$<.001, $\eta_p^2$=.56), condition ($F_{(2,32)}$=5.95, $p$=.006, $\eta_p^2$=.27), and a significant sound x condition interaction ($F_{(2,32)}$=5.96, $p$=.006, $\eta_p^2$=.27). The effect was driven by significant differnces between conditions for target sounds (Down vs. Control $t_{(16)}$=3.62; $p$=.006, d=.62, Down vs. Up $t_{(16)}$=4.57; $p$<.001, $d$=.78, Up vs. Control $t_{(16)}$=0.95; $p$=1), but not for standard sounds (all $p$=1). (d) Scalp ERPs averaged over the time window of interest for the P300 (252-600ms; top) and N100 (50-200ms; bottom). Cluster-based permutation t-tests did not show any significant differences between the modulations conditions or standard and target sounds for the N100. Shaded areas represent s.e.m. Boxplots indicate median (centre line), 25th and 75th percentiles (box), and maximum and minimum values (whiskers). Dots and colours indicate individual participants. Sound icon indicates the presentation of a target or standard sound.
